# Preclinical Comparison of DMT and 5-MeO-DMT Reveals Behavioral Dissociation, Distinct TrkB Activation and Differential Plasticity Profiles

**DOI:** 10.64898/2026.08.06.743248

**Authors:** Orr Shahar, Alexander Botvinnik, Masha Chaykin, Amit Shwartz, Elad Lerer, Peretz Golding, May Ben Ari, Ori Shalev, Tzuri Lifschytz, Bernard Lerer

## Abstract

N, N-dimethyltryptamine (DMT) and 5-methoxy-N, N-dimethyltryptamine (5-MeO-DMT) are structurally related tryptamine psychedelics with emerging therapeutic potential, yet their comparative acute pharmacology and longer-term neuroplastic effects remain incompletely defined. Here we show that DMT produces a bell-shaped dose–response curve in the mouse head-twitch response (HTR) assay, whereas 5-MeO-DMT elicits a monotonic increase. Selective antagonism at 5-HT2A or 5-HT1D receptors, or agonism at 5-HT1A, robustly attenuates HTR for both compounds without abolishing their ability to reduce marble burying, a screening assay for OCD-like behavior. Acutely, both agents elevate TrkB phosphorylation in a region-specific manner, with broader engagement by DMT across default-mode-network and hippocampal territories. Twelve days after a single dose, both compounds increase synaptic proteins (PSD-95, synaptophysin; SV2A for DMT), while DMT uniquely lowers hippocampal BDNF and reprograms frontal-cortex glutathione and energy metabolism. These findings demonstrate that acute hallucinogenic-like activity and selected therapeutic-like behavioral and plasticity outcomes can be pharmacologically dissociated, informing the rational design of more tolerable, scalable psychedelic-based treatments.

## Introduction

Throughout human history, the quest to unravel the enigma of consciousness and the nature of reality has driven our curiosity. It is no surprise, then, that mind-altering substances—often revered and sometimes celebrated —have played a pivotal role in our cultural tapestry. Among these intriguing compounds, N, N-dimethyltryptamine (DMT) stands out. DMT serves as the primary psychedelic compound within the ancient Amazonian brew known as Ayahuasca. Historically, indigenous tribes have employed Ayahuasca for the treatment of physiological and psychological conditions ^1^. DMT is found in various plant species ^2^, and after decades of being hypothesized as endogenously produced in our brains, has recently been shown in rats to be endogenously produced at baseline levels, with a six-fold increase during the physiology of death ^3,4^. Reports suggest that DMT binds to several serotonin receptors (5-HTRs), dopamine receptor 1, adrenergic receptors α1B, α2B, and α2C, and imidazoline receptor 1 ^5,6^. Functional analyses reveal that DMT acts as an agonist at 5-HTRs 1A, 2A, 2B, and 2C ^7,8^, as well as at the rodent trace amine-associated receptor (TAAR 1) ^9^ and the sigma 1 receptor (σ1) ^10^.

A resurgence of therapeutic psychedelic research on DMT, previously stifled by the “war on drugs” during the 1960s, reemerged following clinical trials led by Dr. Rick Strassman in the 1990s, investigating the effects of DMT in humans ^11^. DMT has been a major focus as a potential neuropsychiatric therapeutic agent due to its reported role in synaptic plasticity ^12^. Despite its enigmatic natural role, DMT shares structural similarities with serotonin and melatonin, essential neurotransmitters in the human brain. Its molecular structure differs from serotonin by featuring two methyl groups attached to the nitrogen atom of the tryptamine sidechain, as well as the absence of the 5-hydroxy moiety. Additionally, classic tryptamine psychedelics, such as psilocin (4-OH-DMT), 5-MeO-DMT, and bufotenine (5-OH-DMT)-display motifs from both DMT (two methyl groups attached to the nitrogen atom) and serotonin (an oxygen-based substituent on the indole ring) (Figure 1). While investigating DMT’s natural (intrinsic) function in the brain, we speculate that other tryptamine psychedelics act on shared neural systems involving both serotonin and DMT. These common pathways may explain the profound spiritual experiences reported by users as well as DMT’s documented positive neurobiological effects ^12–14^. One intriguing hypothesis suggests that DMT is released during deep meditative states ^15^. Scientific research on meditation has demonstrated positive physiological ^16^ and neurobiological effects ^17–19^, making it a fascinating avenue for exploring the elusive DMT system. Is the inert DMT system activated through meditation the same system activated when taking an exogenous psychedelic? Is this the mechanism which ancient Amazonian tribes unknowingly relied on when administering Ayahuasca medicinally? Investigation of DMT and Ayahuasca’s therapeutic potential is underway ^20–24^, necessitating further research to elucidate the brain mechanisms activated by DMT specifically and classical psychedelics more generally.

**Figure 1.**
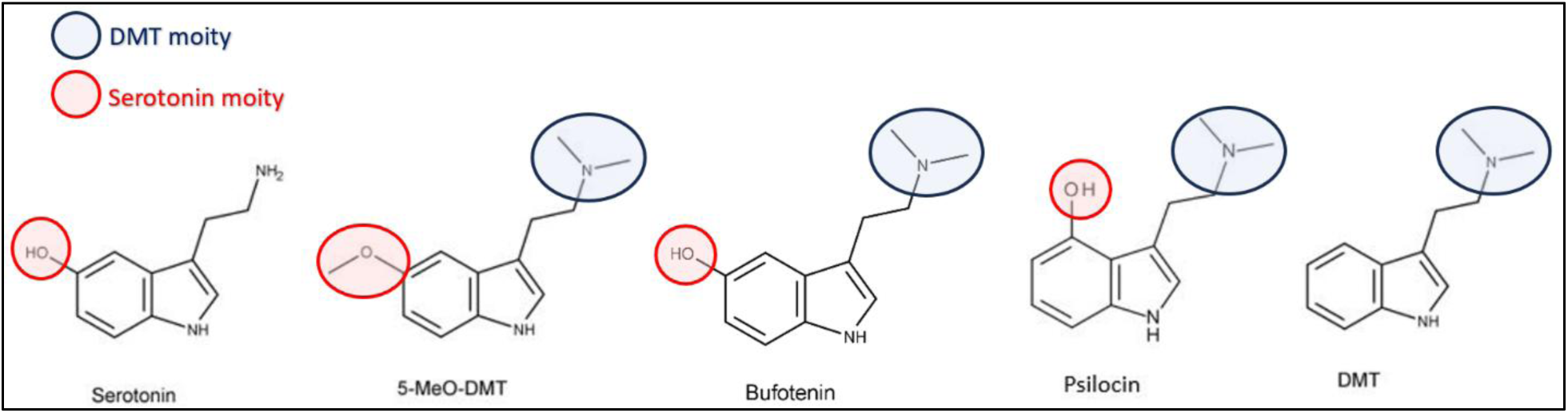
Structural motifs shared by DMT, 5-MeO-DMT and related classic tryptamine psychedelics. Chemical structures of serotonin (5-HT), DMT (N,N-dimethyltryptamine), psilocin (4-OH-DMT), 5-MeO-DMT (5-methoxy-N,N-dimethyltryptamine) and bufotenine (5-OH-DMT). The structures illustrate the common tryptamine core, the characteristic N,N-dimethyl substitution on the ethylamine side chain of DMT-like compounds, and the oxygen-containing substituents at the 4-or 5-position of the indole ring that are present in serotonin and its psychedelic derivatives.

5-Methoxy-N, N-dimethyltryptamine (5-MeO-DMT) is a naturally occurring tryptamine present in various plant species; however, it is best known as the major hallucinogenic agent found in the defensive secretions of *Incilius alvarius* (Colorado River toad) ^25^. Secretions of *Incilius alvarius* have been used by indigenous cultures in the Sonoran Desert region (spanning from southwestern USA to northwestern Mexico) for medicinal and spiritual purposes ^26^. It is a derivative of DMT, which acts as a non-selective ligand at various serotonin receptors, including the G-protein coupled receptors 5-HT1A and 5-HT2A ^27^. 5-MeO-DMT’s high selectivity for 5-HT1A receptors over 5-HT2A receptors suggests that it may differ from typical psychedelics, whose psychoactive effects are currently attributed to 5-HT2A receptor agonism, as demonstrated in human studies ^28,29^. 5-MeO-DMT is also distinct from many other substances that predominantly act as 5-HT1A agonists and have antidepressant and anxiolytic effects ^30^. The psychological and therapeutic potential of 5-MeO-DMT includes alleviating symptoms of depression, anxiety, and post-traumatic stress disorder (PTSD), as well as fostering a sense of interconnection and oneness with the world ^31–34^.

Beyond their acute receptor interactions, a rapidly emerging body of evidence indicates that tryptaminergic psychedelics promote enduring structural and functional neuroplasticity that may underlie their therapeutic potential. While 5-HT2A receptor agonism has long been considered central to the hallucinogenic and many behavioral effects of classic psychedelics, recent work has identified a parallel, 5-HT2A-independent mechanism centered on direct modulation of the BDNF receptor TrkB. In a seminal study, Moliner et al. demonstrated that LSD and psilocin bind directly to the transmembrane domain of TrkB dimers with nanomolar affinity, approximately 1,000-fold higher than that of conventional antidepressants such as fluoxetine or ketamine ^35^. These compounds act as positive allosteric modulators that potentiate and stabilize BDNF-induced TrkB dimerization. LSD exhibits a particularly high residence time on TrkB (k off ≈ 0.0085 min⁻¹), supporting sustained receptor retention on the cell surface for at least 1 h and prolonged downstream signaling (including mTOR phosphorylation). In contrast, BDNF alone typically elicits transient TrkB dimerization and signaling on the scale of minutes. This allosteric facilitation provides a molecular substrate for the rapid and persistent neuroplasticity observed after psychedelic administration.

Importantly, the plasticity-promoting effects of psychedelics (TrkB dimerization, spinogenesis, dendritogenesis, and antidepressant-like behaviors) persist in the presence of 5-HT2A antagonists and are abolished by TrkB-binding-site mutations or BDNF sequestration, whereas the head-twitch response (HTR) (a well-validated rodent proxy for hallucinogenic potential) remains strictly 5-HT2A-dependent and TrkB-independent ^35^. This mechanistic dissociation further supports the possibility that the intense subjective effects and the long-term therapeutic plasticity may be pharmacologically separable.

In our recent work comparing chemically synthesized psilocybin (PSIL) with a psychedelic mushroom extract (PME), we observed that PME produced more potent and prolonged increases in synaptic plasticity markers together with distinct frontal-cortex metabolomic signatures ^36^. These findings prompted us to ask whether single-dose administration of the structurally related tryptamines DMT and 5-MeO-DMT would similarly induce lasting molecular and metabolic changes. Accordingly, the present study examined both acute (4 min post-injection, corresponding to peak HTR) and long-term (12 days) cortical metabolomic profiles, alongside hippocampal BDNF/TrkB protein levels and expression of the synaptic-plasticity-related proteins GAP43, PSD95, synaptophysin, and SV2A at the 12-day time point. We further quantified region-specific (default mode network and hippocampus) TrkB phosphorylation (p-TrkB/TrkB immunofluorescence) 1 h after treatment to capture early activation of this plasticity pathway.

A second, complementary arm of the study addressed the translational goal of developing more accessible psychedelic-based treatments. Because the profound acute subjective effects of psychedelics can limit tolerability and scalability, we systematically tested whether selective agonists or antagonists at key serotonergic receptors (5-HT1A, 5-HT1B, 5-HT1D, 5-HT2A, 5-HT2C, 5-HT7) and other targets (TAAR1, σ1) could attenuate the HTR while preserving therapeutic-like behavioral effects. For the latter, we employed the marble-burying test (MBT), a widely used screening assay for compounds that alleviate OCD-like symptoms. Notably, 5-HT1A receptor agonists such as buspirone reduce marble burying, and this behavioral output has been shown to be sensitive to several serotonergic psychedelics ^37^. We therefore asked whether co-administration of receptor modulators that blunt HTR would also blunt, or conversely spare, the reduction in marble burying produced by DMT or 5-MeO-DMT. Preservation of the MBT effect despite HTR attenuation would support the feasibility of pharmacological strategies that reduce acute hallucinogenic-like activity without sacrificing potential therapeutic benefit in this screening test for OCD-like symptom alleviation.

By integrating acute pharmacological dissection of HTR and MBT with parallel molecular profiling of TrkB activation, long-term synaptic protein expression, and metabolomic reprogramming, the present work provides a comprehensive preclinical characterization of DMT and 5-MeO-DMT. The findings are intended to inform both mechanistic understanding of DMT based-induced plasticity and the rational design of combination regimens that could help develop psychedelic-constructed therapeutic tools that are more tolerable and broadly accessible.

## Results

### Head Twitch Response

To compare the effects of DMT and 5-MeO-DMT, we first characterized their dose-response relationships on the head twitch response (HTR) across the same dose range (1.25–80 mg/kg, i.p.) (Figure 2). One-way ANOVA revealed a significant overall effect of DMT dose on total HTR (F (7, 41) = 10.85, P < 0.001). Post hoc Dunnett’s multiple comparisons tests showed that DMT significantly increased total HTR at 1.25 mg/kg (p = 0.001), 2.5 mg/kg (p < 0.001), and 5 mg/kg (p = 0.003) compared to vehicle. Higher doses (10– 80 mg/kg) did not induce a significant increase in HTR relative to vehicle (Figure 2a, b).

**Figure 2.**
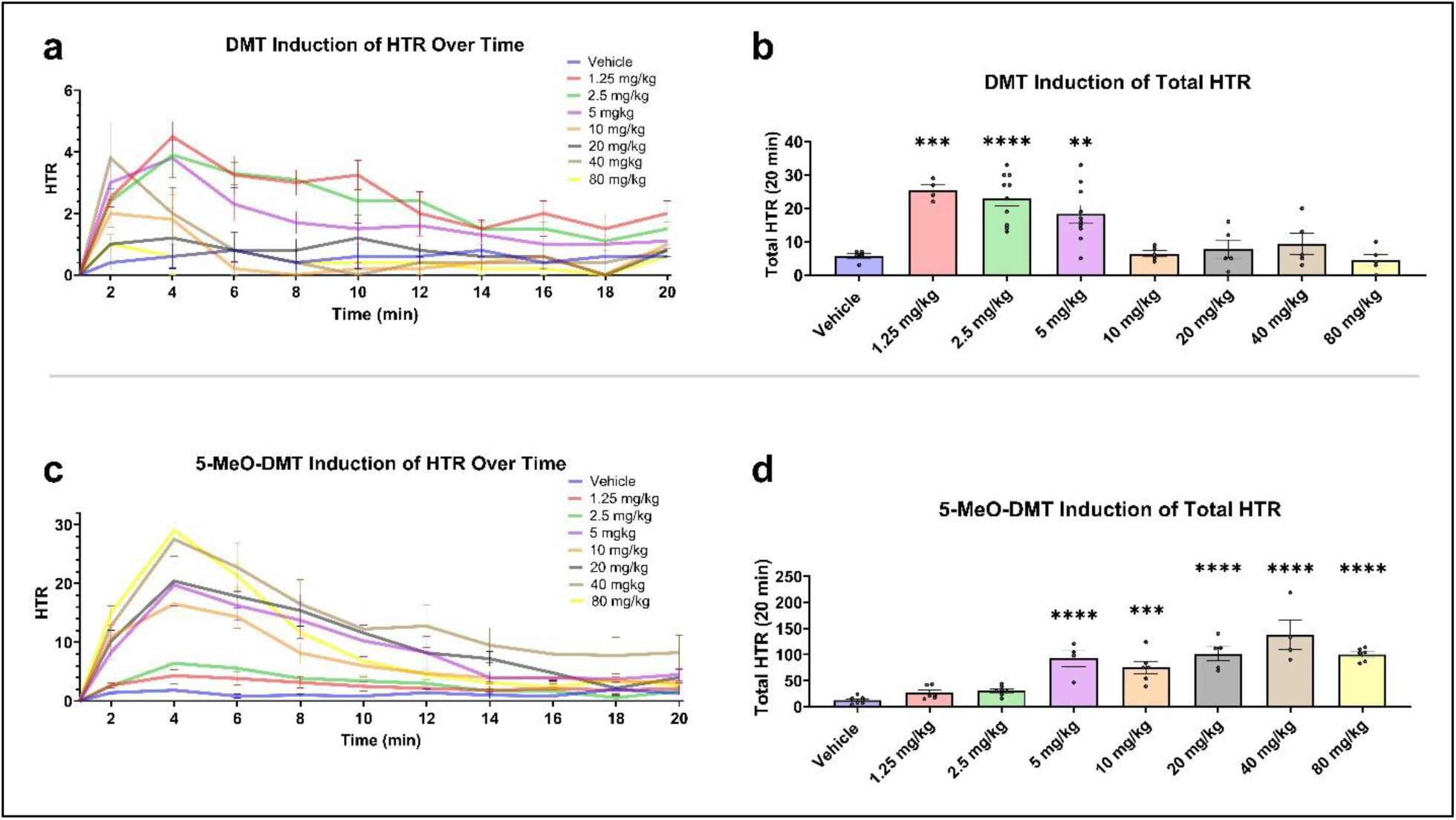
DMT induces a bell-shaped dose-response profile in the mouse head-twitch response (HTR) assay, whereas 5-MeO-DMT induces a monotonic dose-response relationship. (a) Head twitches recorded over 20 min (presented in two-min time-bins) following i.p. administration of vehicle (VEH) or DMT (1.25–80 mg/kg). (b) Total HTR induced by each DMT dose or VEH during the 20-min session. One-way ANOVA: F(7, 41) = 10.85, P < 0.001. Dunnett’s post-hoc tests vs vehicle: 1.25 mg/kg and 2.5 mg/kg P < 0.001; 5 mg/kg P = 0.004; 10–80 mg/kg P > 0.05. (c) Head twitches recorded over 20 min (presented in two-min time-bins) following i.p. administration of vehicle (VEH) or 5-MeO-DMT (1.25–80 mg/kg). (d) Total HTR induced by each 5-MeO-DMT dose or VEH during the 20-min session. One-way ANOVA: F(7, 37) = 17.47, P < 0.001. Dunnett’s post-hoc tests vs vehicle: 1.25 and 2.5 mg/kg P > 0.05; 5–80 mg/kg P < 0.001. Data are mean ± SEM HTR in 2 min time bins, over 20 min. n = 4–10 (a, b) or 4–7 (c, d). ** P < 0.01, *** P < 0.001, **** P < 0.0001.

One-way ANOVA revealed a significant overall effect of 5-MeO-DMT dose on total HTR (F (7, 37) = 17.47, P < 0.001). Post hoc Dunnett’s multiple comparisons tests showed that 5-MeO-DMT significantly increased HTR compared to vehicle at the doses of 5 mg/kg (p < 0.001), 10 mg/kg (p = 0.001), 20 mg/kg (p < 0.001), 40 mg/kg (p < 0.001), and 80 mg/kg (p < 0.001) (Figure 2c, d).

Doses for subsequent experiments were selected based on translational relevance, calculated using the DoseCal tool ^38^, and observed HTR efficacy. We used 5 mg/kg DMT, as it produced significant HTR and is translationally relevant. For 5-MeO-DMT, we used 10 mg/kg in pharmacological and molecular studies to ensure a clear supra-threshold dose that elicits strong and reliable HTR, well above the lower end of the effective dose range observed in our assays. In behavioral experiments, we used the lower dose of 5 mg/kg to maintain translational relevance.

### Pharmacological modulation of HTR

To investigate receptor mechanisms underlying DMT-and 5-MeO-DMT-induced HTR, we tested the effects of selective agonists and antagonists at multiple serotonin receptor subtypes and the sigma receptor (Figure 3). One-way ANOVA revealed a significant overall effect in the DMT pharmacological modulation experiment (F (11, 50) = 7.544, P < 0.001). Post hoc Dunnett’s tests showed that the 5-HT2A antagonist M107900 (0.5 mg/kg) significantly attenuated DMT-induced HTR (p = 0.029). The 5-HT1A agonist 8-OHDPAT (1 mg/kg) also significantly reduced DMT-induced HTR (p = 0.038), whereas the 5-HT1A antagonist NAD-299 (3 mg/kg) significantly increased DMT-induced HTR (p < 0.001). The 5-HT1D antagonist LY310762 significantly reduced DMT-induced HTR (p = 0.027). Co-administration with the 5-HT1D agonist PNU 142633 (0.3 mg/kg), 5-HT2C agonist WAY-161503 (3 mg/kg), 5-HT2C antagonist RS-102221 (4 mg/kg), 5-HT7 agonist AS-19 (20 mg/kg), 5-HT7 antagonist SB-269970 (60mg/kg), TAAR1 antagonist EPPTB (10mg/kg), and σ receptor antagonist BD-1047 (10 mg/kg) did not significantly affect DMT-induced HTR (Figure 3a).

**Figure 3.**
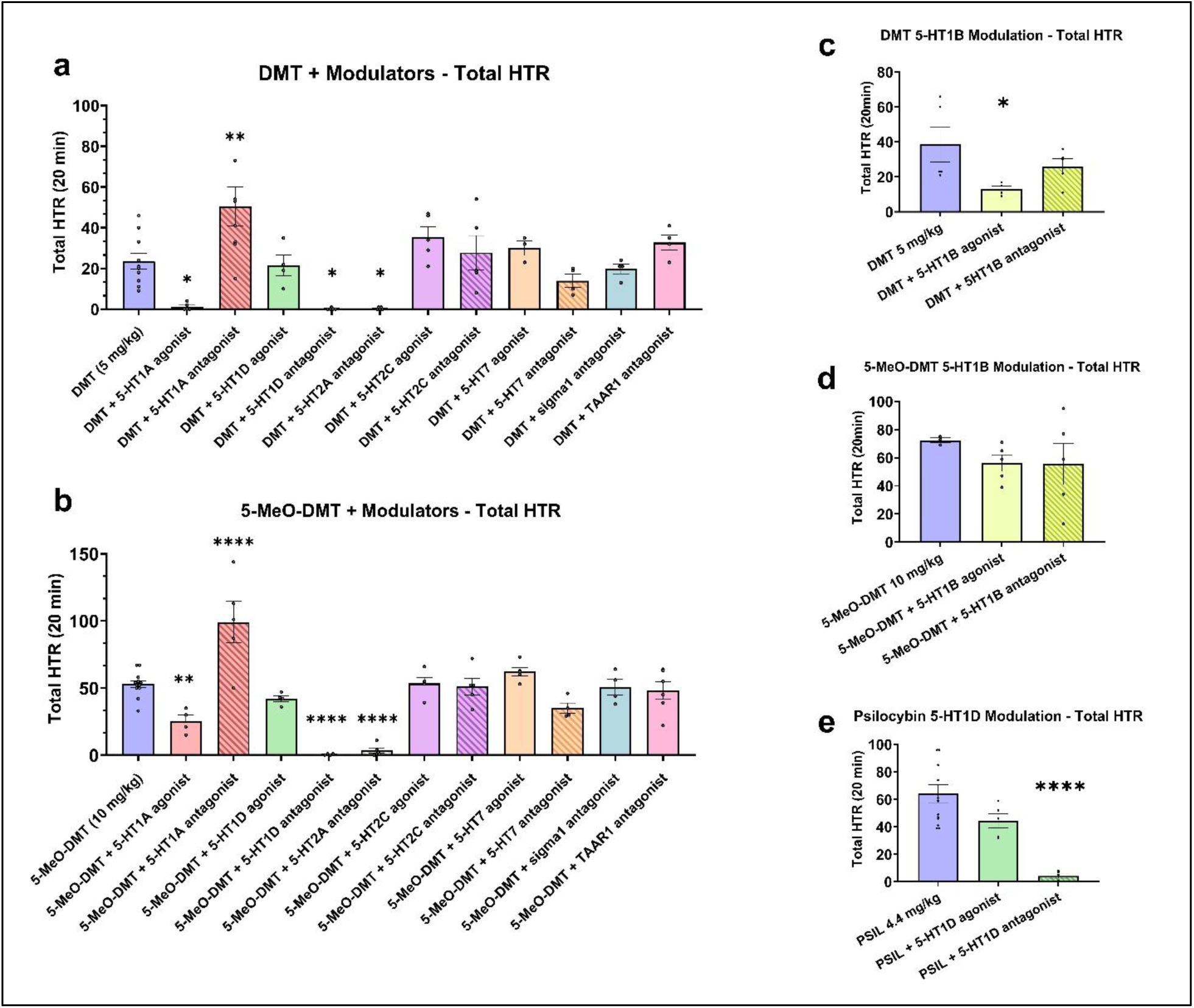
Pharmacological receptor modulation of DMT-and 5-MeO-DMT-induced head-twitch responses, with comparable psilocybin-induced HTR 5-HT1D receptor modulation. (a) Effect of selective agonists and antagonists on DMT (5 mg/kg)-induced HTR. One-way ANOVA: F(11, 50) = 7.544, P < 0.001. Dunnett’s post-hoc tests vs DMT alone: 5-HT2A antagonist M107900 (0.5 mg/kg) P = 0.029; 5-HT1A agonist 8-OH-DPAT (1 mg/kg) P = 0.039; 5-HT1A antagonist NAD-299 (3 mg/kg) P = 0.001; 5-HT1D antagonist LY310762 (1 mg/kg) P = 0.027; all other modulators P > 0.05. (b) Effect of the same modulators on 5-MeO-DMT (10 mg/kg)-induced HTR. One-way ANOVA: F(11, 52) = 18.25, P < 0.001. Dunnett’s post-hoc tests vs 5-MeO-DMT alone: 5-HT2A antagonist M107900 (0.5 mg/kg), 5-HT1A antagonist NAD-299 (3 mg/kg), and 5-HT1D antagonist LY310762 (1 mg/kg) P < 0.001; 5-HT1A agonist 8-OH-DPAT (1 mg/kg) P = 0.007; all other modulators P > 0.05. (c) Focused analysis of DMT (5 mg/kg) ± 5-HT1B receptor modulation. One-way ANOVA: F(2, 11) = 4.86, P = 0.031. Dunnett’s post-hoc tests vs DMT alone: 5-HT1B agonist CP 94253 (5 mg/kg) P = 0.019; 5-HT1B antagonist SB 224289 (2.5 mg/kg) P = 0.44. (d) Focused analysis of 5-MeO-DMT (10 mg/kg) ± 5-HT1B receptor modulation. Neither the 5-HT1B agonist CP 94253 (5 mg/kg) nor the 5-HT1B antagonist SB 224289 (2.5 mg/kg) significantly modulated 5-MeO-DMT-induced HTR (both P > 0.05). (e) Focused analysis of PSIL (psilocybin) (4.4 mg/kg) ± 5-HT1D receptor modulation. One-way ANOVA: F(2, 20) = 51.41, P < 0.001. Dunnett’s post-hoc tests vs PSIL alone: 5-HT1D agonist PNU 142633 (0.3 mg/kg) P = 0.417, 5-HT1D antagonist LY310762 (1 mg/kg) P < 0.001. Data are mean ± SEM total HTR in 20 min. One-way ANOVA followed by Dunnett’s post-hoc test. * P < 0.05, ** P < 0.01, *** P < 0.001, **** P < 0.0001. n = 4–10 (a, b) or n = 3–5 (c, d).

One-way ANOVA revealed a significant overall effect in the 5-MeO-DMT pharmacological modulation experiment (F (11, 52) = 18.25, P < 0.001). Post hoc Dunnett’s tests showed that the 5-HT2A antagonist M107900 (0.5 mg/kg) significantly attenuated 5-MeO-DMT-induced HTR (p < 0.001). The 5-HT1A agonist 8-OHDPAT (1 mg/kg) also significantly reduced 5-MeO-DMT-induced HTR (p = 0.007), while the 5-HT1A antagonist NAD-299 (3 mg/kg) significantly increased 5-MeO-DMT-induced HTR (p < 0.001). The 5-HT1D antagonist LY310762 (1 mg/kg) also significantly decreased 5-MeO-DMT-induced HTR (p < 0.001). Co-administration with the 5-HT1D agonist PNU 142633 (0.3 mg/kg), 5-HT2C agonist WAY-161503 (3 mg/kg), 5-HT2C antagonist RS-102221 (4 mg/kg), 5-HT7 agonist AS-19 (20 mg/kg), 5-HT7 antagonist SB-269970 (60mg/kg), TAAR1 antagonist EPPTB (10mg/kg), and σ receptor antagonist BD-1047 (10 mg/kg) did not significantly affect 5-MeO-DMT-induced HTR (Figure 3b).

The effects of the 5-HT1B receptor were examined in separate focused experiments using one-way ANOVA (Figure 3c, d). For DMT, one group showed a borderline deviation from normality on the log transform of the data. Given the small sample sizes and consistency with previous DMT HTR experiments, parametric analysis was performed on the log transformed data. One-way ANOVA revealed a significant overall effect (F (2, 11) = 4.86, P = 0.031). Post hoc Dunnett’s tests showed that co-administration of the 5-HT1B agonist CP 94253 (5 mg/kg) significantly reduced total HTR induced by DMT (p = 0.019). In contrast, co-administration of the 5-HT1B antagonist SB 224289 (2.5 mg/kg) did not significantly affect DMT-induced HTR. Neither the 5-HT1B agonist CP 94253 (5 mg/kg) nor the 5-HT1B antagonist SB 224289 (2.5 mg/kg) significantly modulated 5-MeO-DMT-induced HTR.

As an addition to our previously reported psilocybin-induced HTR modulation ^39^, we comparably assessed the effects of 5-HT1D receptor modulation on psilocybin-induced HTR in a separate focused experiment (Figure 4e). In this experiment the datasets met normality assumptions after log transformation, therefore parametric analysis was performed on the log transformed data. The one-way ANOVA test revealed a significant overall effect (F (2, 20) = 51.41, P < 0.001). Post hoc Dunnett’s tests showed that co-administration of the 5-HT1D antagonist LY310762 (1 mg/kg) significantly reduced total HTR induced by DMT (P < 0.001). In contrast, co-administration of the 5-HT1D agonist PNU 142633 (0.3 mg/kg) did not significantly affect psilocybin-induced HTR.

**Figure 4.**
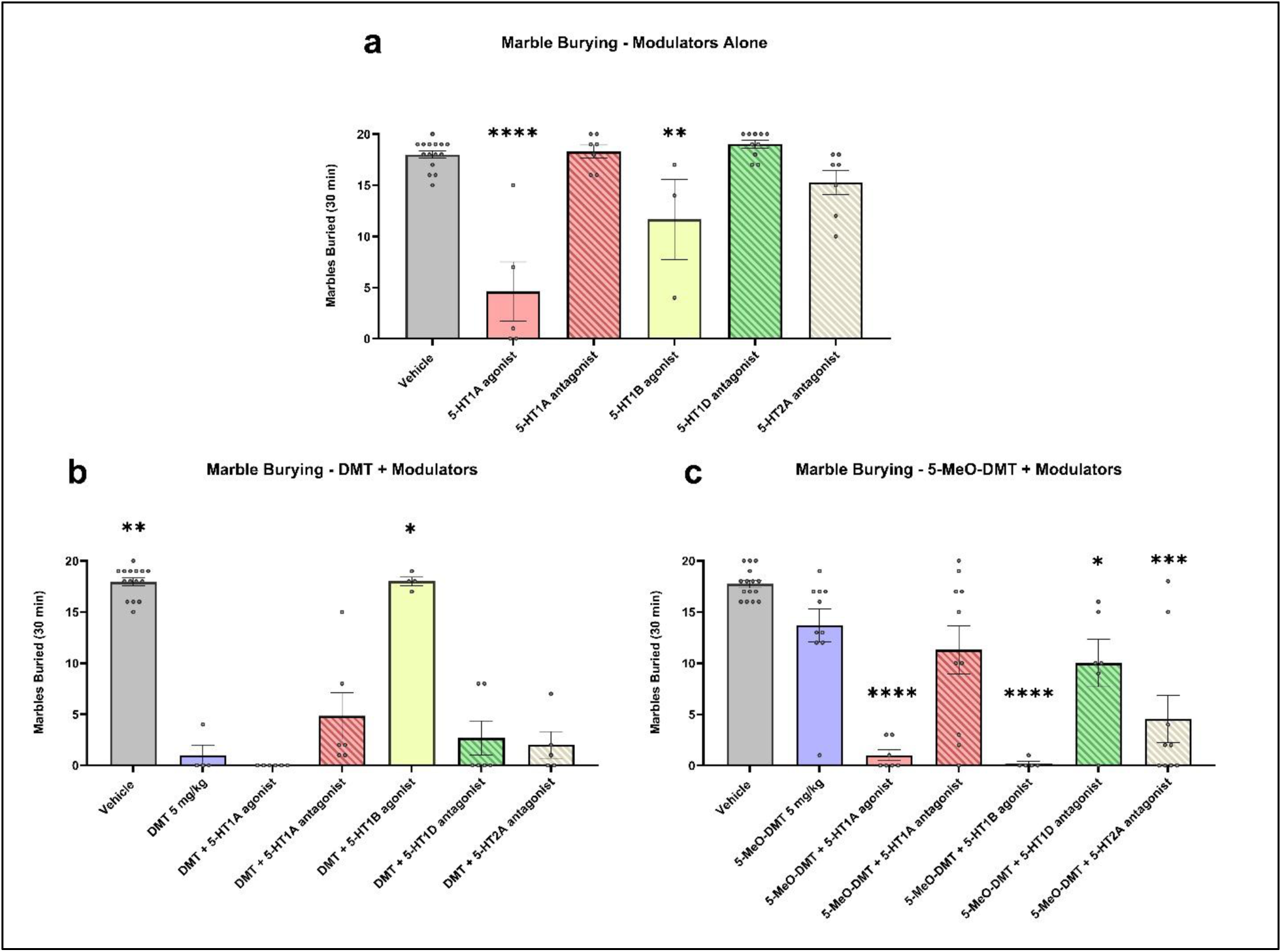
Effects of receptor modulators, DMT, and 5-MeO-DMT on marble-burying behavior. Data show the number of marbles buried during a 30-min test session. (a) Effects of receptor modulators administered alone. One-way ANOVA: F(5, 41) = 19.30, P < 0.001. Dunnett’s post-hoc tests vs vehicle: 5-HT1A agonist 8-OH-DPAT (1 mg/kg) P < 0.001; 5-HT1B agonist CP 94253 (5 mg/kg) P = 0.010; 5-HT2A antagonist M107900 (0.5 mg/kg), 5-HT1A antagonist NAD-299 (3 mg/kg), and 5-HT1D antagonist LY310762 (1 mg/kg) P > 0.05. (b) Effects of DMT (5 mg/kg) ± receptor modulators. Kruskal–Wallis test: P < 0.001. Dunn’s post-hoc tests vs DMT: vehicle P = 0.003; DMT + 5-HT1B agonist CP 94253 (5 mg/kg) P = 0.046; DMT + 5-HT2A antagonist M107900 (0.5 mg/kg), DMT + 5-HT1A agonist 8-OH-DPAT (1 mg/kg), DMT + 5-HT1A antagonist NAD-299 (3 mg/kg), and DMT + 5-HT1D antagonist LY310762 (1 mg/kg) P > 0.05. (c) Effects of 5-MeO-DMT (5 mg/kg) ± receptor modulators. Kruskal–Wallis test: P < 0.001. Dunn’s post-hoc tests vs vehicle: 5-MeO-DMT P = 0.605; 5-MeO-DMT + 5-HT2A antagonist M107900 (0.5 mg/kg), 5-MeO-DMT + 5-HT1A agonist 8-OH-DPAT (1 mg/kg), and 5-MeO-DMT + 5-HT1B agonist CP 94253 (5 mg/kg) P < 0.001. Data are mean ± SEM total marbles buried in 30 min. * P < 0.05, ** P < 0.01, *** P < 0.001, **** P < 0.0001.

A summary of the pharmacological modulation of HTR induced by DMT, 5-MeO-DMT, and as previously presented by psilocybin ^39^ with the addition of 5-HT1D modulation is presented in Table 1.

**Table 1.**
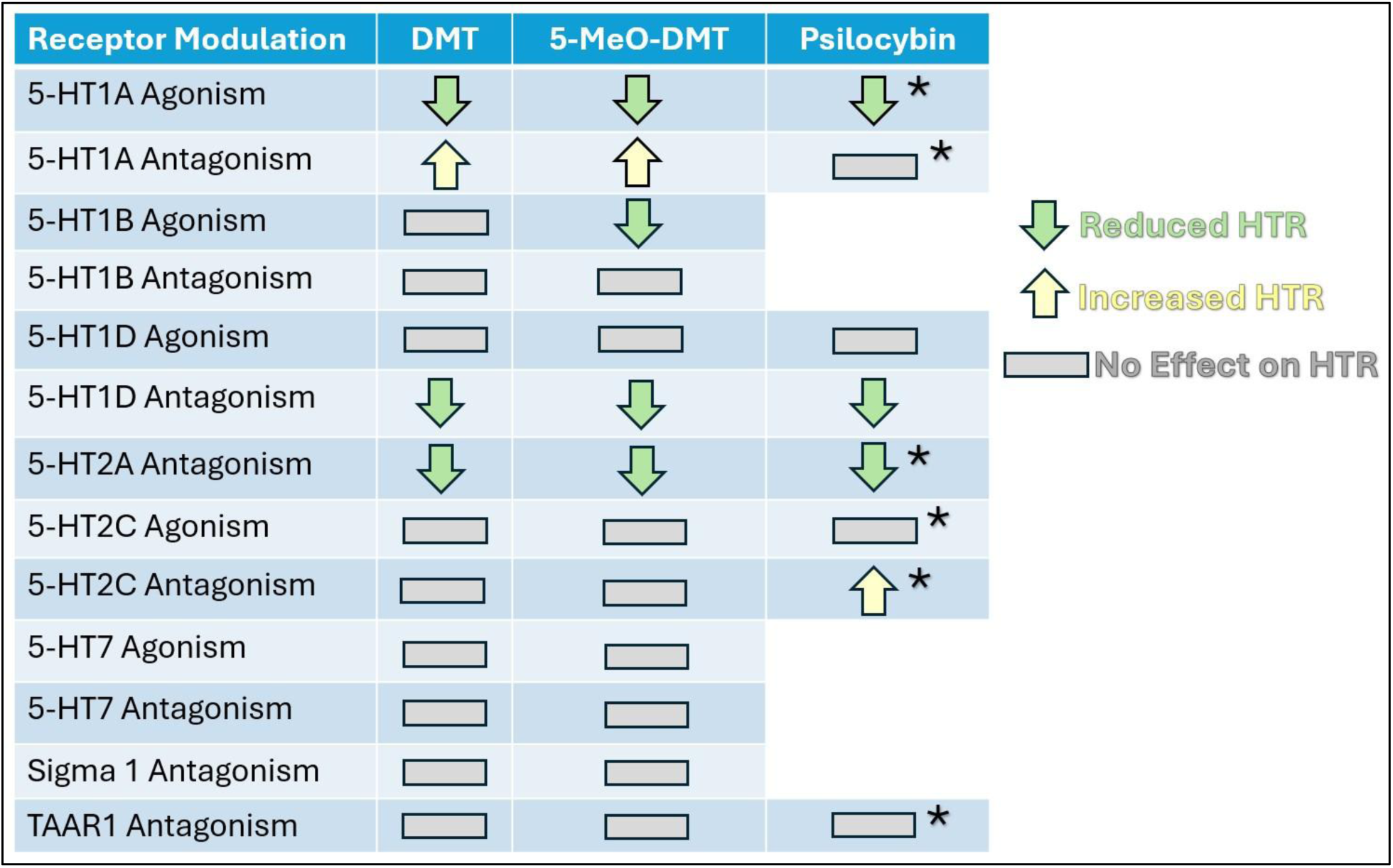
Summary of a specific receptor modulation on the HTR induced by DMT, 5-MeO-DMT, and psilocybin treatments. Based on the data collected from Figure 3, and adding onto data previously reported ^39^. An Arrow up represents an observed increase of HTR (P < 0.05), an arrow down represents a statistically significant reduction of HTR (P < 0.05), and a rectangle represents no effect on the HTR induced by DMT, 5-MeO-DMT, and psilocybin treatments. Astrix (*) represents data taken from our previously reported study and is used as a comparable reference only and not as a direct result from the current study.

### Marble Burying

We examined whether receptor modulators that affected the head-twitch response also influenced marble burying behavior when administered alone or in combination with DMT and 5-MeO-DMT (Figure 4).

First, we tested the modulators alone. One-way ANOVA revealed a significant overall effect (F (5, 41) = 19.30, P < 0.001). Post hoc Dunnett’s multiple comparisons tests showed that the 5-HT1A agonist 8-OHDPAT (1 mg/kg) and the 5-HT1B agonist CP 94253 (5 mg/kg) significantly reduced marble burying compared to vehicle (p < 0.001 and P = 0.01, respectively). In contrast, the 5-HT2A antagonist M107900 (0.5 mg/kg), the 5-HT1A antagonist NAD-299 (3 mg/kg), and the 5-HT1D antagonist LY310762 (1 mg/kg) did not significantly affect marble burying on their own.

Because the DMT and 5-MeO-DMT datasets did not meet normality assumptions even after log transformation, non-parametric analyses were used. In the DMT experiment, the Kruskal–Wallis test revealed a significant overall effect (p < 0.001). Dunn’s post-hoc tests showed that DMT (5 mg/kg) alone significantly reduced marble burying compared to vehicle (p = 0.003). This reduction was significantly attenuated by co-administration of the 5-HT1B agonist CP 94253 (5 mg/kg) (DMT vs DMT + 5-HT1B agonist, P = 0.045). Co-administration with the 5-HT2A antagonist M107900 (0.5 mg/kg), 5-HT1A agonist 8-OHDPAT (1 mg/kg), 5-HT1A antagonist NAD-299 (3 mg/kg), or 5-HT1D antagonist LY310762 (1 mg/kg) did not significantly alter the effect of DMT.

In the 5-MeO-DMT experiment, the Kruskal–Wallis test also revealed a significant overall effect (p < 0.001). Because 5-MeO-DMT (5 mg/kg) alone did not significantly reduce marble burying compared to vehicle, all groups were compared to vehicle to assess effects of administration with the co-treatments. Dunn’s post-hoc tests showed that co-administration of 5-MeO-DMT with the 5-HT2A antagonist M107900 (0.5 mg/kg), the 5-HT1A agonist 8-OHDPAT (1 mg/kg), or the 5-HT1B agonist CP 94253 (5 mg/kg) significantly reduced marble burying compared to vehicle (all P < 0.001). Notably, co-administration of 5-MeO-DMT with the 5-HT1B agonist CP 94253 (5 mg/kg) was associated with mortality in some animals.

### TrkB activation by DMT and 5-MeO-DMT

Acute administration of DMT (5 mg/kg) or 5-MeO-DMT (5 mg/kg) increased TrkB phosphorylation (p-TrkB/TrkB ratio) in mouse brain, as assessed by immunofluorescence 1 h after treatment (Figure 5a). Data were log-transformed and analyzed by two-way ANOVA, which revealed a significant main effect of treatment (F (2, 114) = 36.69, P < 0.001) and brain region (F (10, 114) = 4.932, P < 0.001), with no significant interaction. Post hoc Tukey’s multiple comparisons tests showed that DMT significantly increased TrkB phosphorylation compared to vehicle in default mode network areas including the frontal association cortex (FrA), prelimbic cortex (PrL), medial orbital/infralimbic cortex (MO-IL), secondary motor cortex (M2), retrosplenial dysgranular cortex (RSD), and retrosplenial granular cortex (RSG). DMT also significantly elevated TrkB phosphorylation in hippocampal regions including CA1, CA2, and the dentate gyrus (DG) (all P < 0.05), but not in the cingulate cortex areas Cg1 and Cg2. In contrast, 5-MeO-DMT produced significant increases only in the FrA and PrL (p < 0.05). Direct comparison between the two compounds revealed that DMT induced significantly greater TrkB phosphorylation than 5-MeO-DMT in the CA2 (P < 0.05) region and DG (P = 0.004).

**Figure 5.**
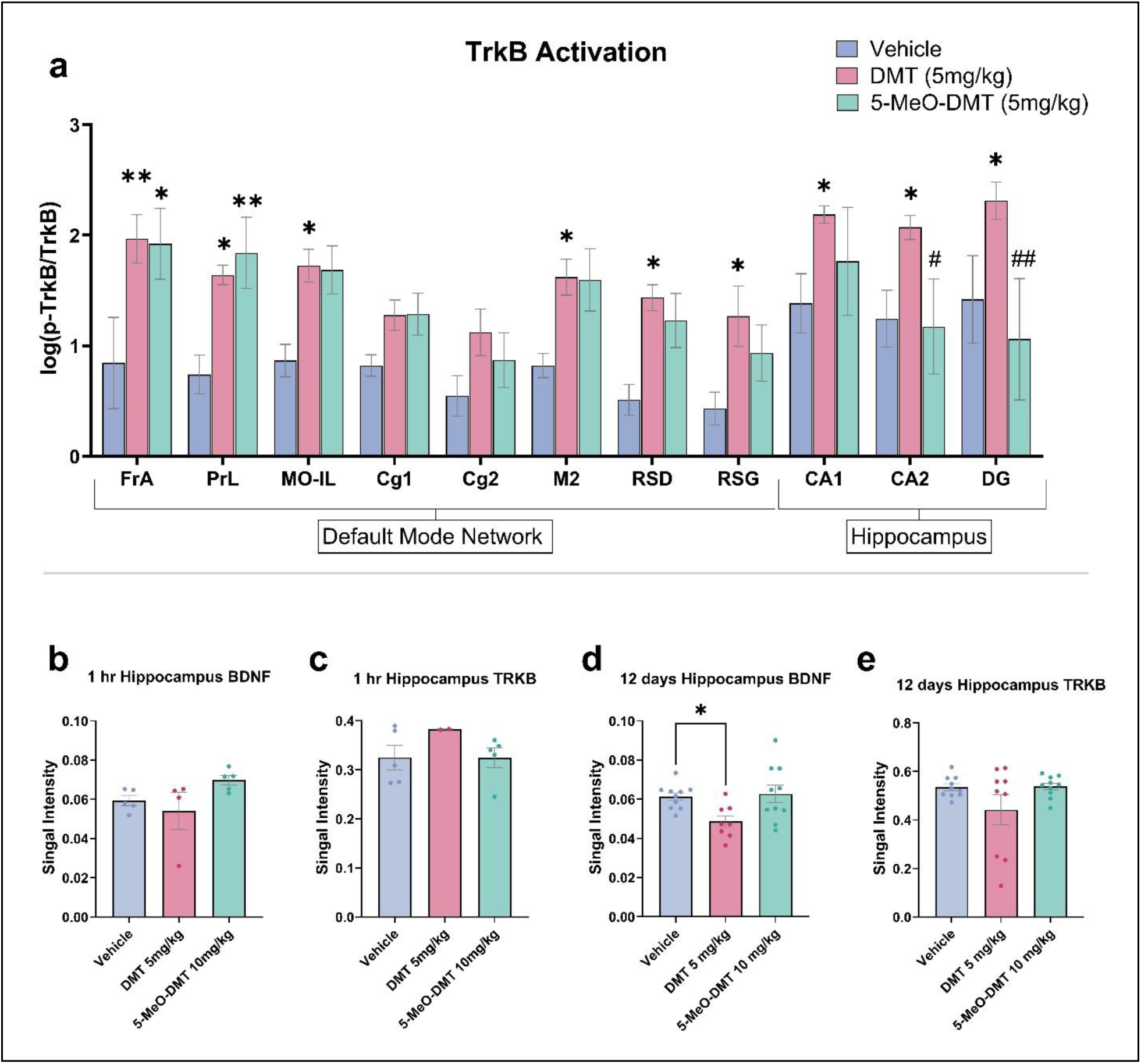
TrkB activation and BDNF/TrkB protein levels following DMT and 5-MeO-DMT administration. (a) Acute TrkB phosphorylation (p-TrkB / unphosphorylated TrkB ratio) in mouse brain regions 1 h after administration of DMT (5 mg/kg) or 5-MeO-DMT (5 mg/kg). Data were log-transformed and analyzed by two-way ANOVA: main effect of treatment F(2, 114) = 36.69, P < 0.001; main effect of brain region F(10, 114) = 4.932, P < 0.001; no significant interaction. Post-hoc Tukey’s tests: DMT significantly increased p-TrkB/TrkB vs vehicle in FrA, PrL, MO-IL, M2, RSD, RSG, CA1, CA2, and DG (all P < 0.05), but not in Cg1 or Cg2. 5-MeO-DMT significantly increased p-TrkB/TrkB only in FrA and PrL (P < 0.05). DMT induced significantly greater phosphorylation than 5-MeO-DMT in CA2 (P < 0.05) and DG (P = 0.004). (b) Hippocampal BDNF protein levels 1 h after administration. Kruskal–Wallis test: P = 0.037. Dunn’s post-hoc: Vehicle vs DMT P > 0.999; Vehicle vs 5-MeO-DMT P = 0.070; DMT vs 5-MeO-DMT P = 0.156. (c) Hippocampal TrkB protein levels 1 h after administration. Kruskal–Wallis test: P = 0.243 (no significant differences). (d) Hippocampal BDNF protein levels 12 days after administration. One-way ANOVA: F(2, 25) = 4.751, P = 0.018. Dunnett’s post-hoc: DMT vs vehicle P = 0.031; 5-MeO-DMT vs vehicle P = 0.939. (e) Hippocampal TrkB protein levels 12 days after administration. One-way ANOVA: F(2, 26) = 2.328, P = 0.118 (no significant differences). Data are mean ± SEM. * P < 0.05, ** P < 0.01 vs vehicle. # P < 0.05, ## P < 0.01 vs DMT. n = 3–5 (a) or 4–10 (b–e).

### Analysis of BDNF and TrkB Proteins following DMT and 5-MeO-DMT administration

The expression levels of BDNF and TrkB proteins in the hippocampus were assessed 1 hour and 12 days after administration of 5 mg/kg DMT or 10 mg/kg 5-MeO-DMT and compared to the vehicle control group. At 1 hour after administration, BDNF levels showed a significant overall difference among groups (Kruskal– Wallis test, P = 0.037); however, Dunn’s post-hoc multiple comparisons showed no statistically significant pairwise differences (Figure 5b). Acute administration of DMT (5 mg/kg) or 5-MeO-DMT (10 mg/kg) did not significantly alter TrkB protein levels in the hippocampus 1 h after injection (Figure 5c). Non-parametric tests were used for the 1-hour data because both the raw and log-transformed values failed normality assumptions.

Twelve days after administration, BDNF levels showed a significant overall difference (one-way ANOVA, F (2, 25) = 4.751, P = 0.017) (Figure 5d). Dunnett’s post-hoc test indicated that DMT significantly reduced BDNF expression compared with vehicle (p = 0.031), whereas 5-MeO-DMT did not induce a significant change compared to vehicle. In contrast, hippocampal TrkB protein levels remained unchanged across treatment groups (Figure 5e).

### Effect of DMT and 5-MeO-DMT on synaptic proteins

To assess long-term effects on synaptic plasticity, levels of GAP43, PSD95, synaptophysin, and SV2A were measured by Western blot in homogenates from the frontal cortex, amygdala, and hippocampus 12 days after treatment (Figure 6). Nested one-way ANOVA revealed no significant overall effect of treatment on GAP43 levels across brain regions.

**Figure 6.**
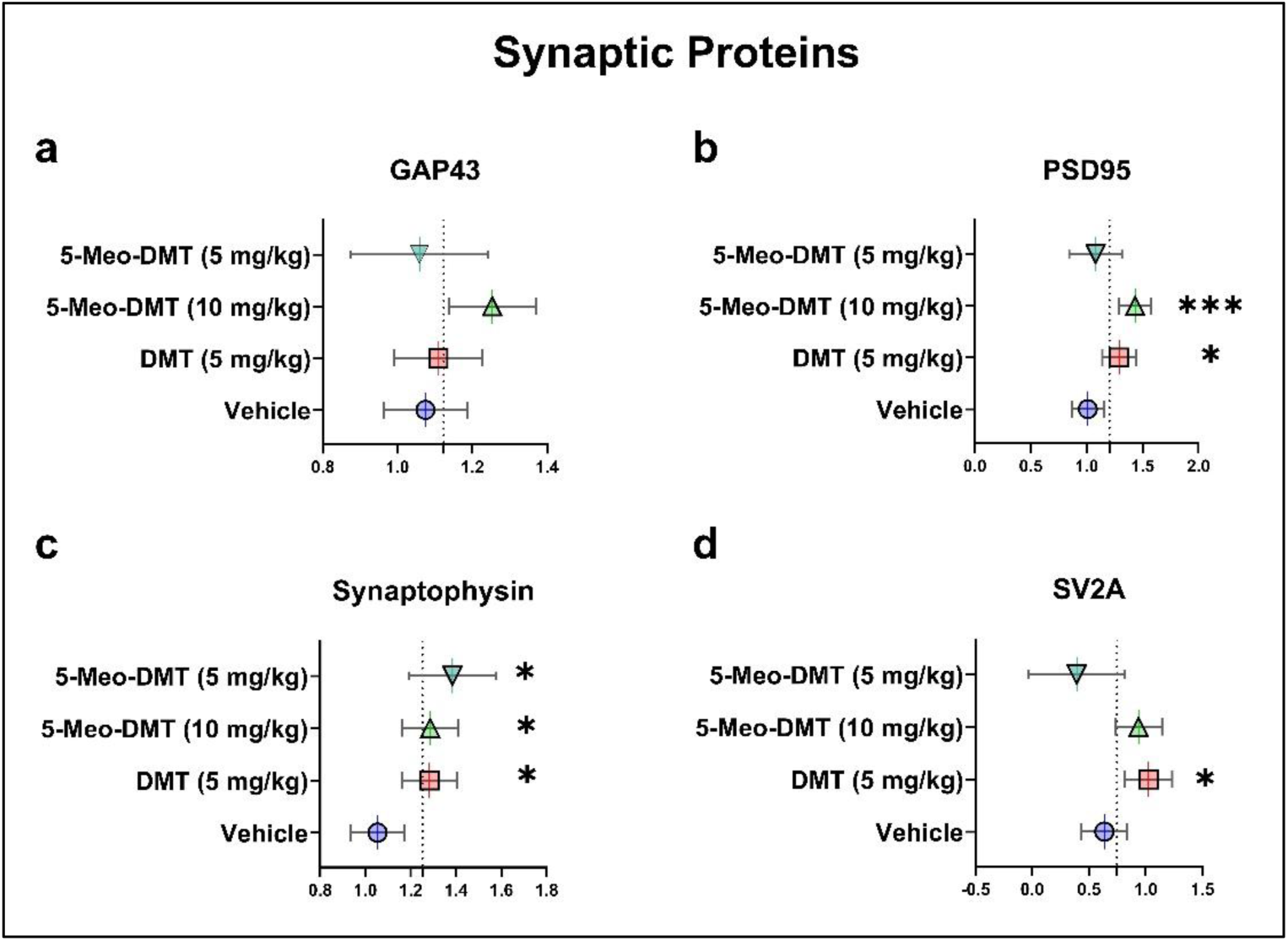
Long-term effects of a single dose of DMT or 5-MeO-DMT on synaptic plasticity-related proteins. Levels of GAP43, PSD95, synaptophysin, and SV2A were quantified by Western blot in the frontal cortex, amygdala, and hippocampus 12 days after treatment. Nested one-way ANOVA (brain region nested within treatment) followed by Dunnett’s post-hoc test. (a) GAP43: Nested one-way ANOVA: F(3, 142) = 2.005, P = 0.116 (no significant overall effect). (b) PSD95: Nested one-way ANOVA: F(3, 147) = 6.335, P < 0.001. Dunnett’s post-hoc vs vehicle: DMT (5 mg/kg) P = 0.024; 5-MeO-DMT (10 mg/kg) P < 0.001; 5-MeO-DMT (5 mg/kg) P > 0.05. (c) Synaptophysin: Nested one-way ANOVA: F(3, 149) = 4.250, P = 0.007. Dunnett’s post-hoc vs vehicle: DMT (5 mg/kg), 5-MeO-DMT (10 mg/kg), and 5-MeO-DMT (5 mg/kg) P < 0.05. (d) SV2A: Nested one-way ANOVA: F(3, 137) = 4.128, P = 0.008. Dunnett’s post-hoc vs vehicle: DMT (5 mg/kg) P = 0.024; both doses of 5-MeO-DMT P > 0.05. Data are mean ± SEM normalized to β-actin. * P < 0.05, *** P < 0.001. n = 5–16 per group. Nested analyses for individual brain regions are provided in Supplementary Figure 1.

For PSD95, nested one-way ANOVA showed a significant overall effect of treatment (F (3, 147) = 6.335, P < 0.001). Post hoc Dunnett’s tests indicated significant increases compared to vehicle following DMT (5 mg/kg) (p = 0.023) and 5-MeO-DMT (10 mg/kg) (p < 0.001), but not following 5-MeO-DMT (5 mg/kg).

For synaptophysin, nested one-way ANOVA revealed a significant overall effect of treatment (F (3, 149) = 4.250, P = 0.006). Post hoc Dunnett’s tests showed significant increases compared to vehicle following DMT (5 mg/kg) (p = 0.024), 5-MeO-DMT (10 mg/kg) (p = 0.021), and 5-MeO-DMT (5 mg/kg) (p = 0.012).

For SV2A, nested one-way ANOVA showed a significant overall effect of treatment (F (3, 137) = 4.128, P = 0.007). Post hoc Dunnett’s tests indicated a significant increase compared to vehicle following DMT treatment (5 mg/kg) (p = 0.023), but not following either dose of 5-MeO-DMT. Individual nested one-way ANOVA results for each brain region are provided in Supplementary Figure 1.

### Metabolomics

Untargeted liquid chromatography–mass spectrometry (LCMS)-based metabolomics was performed on mouse frontal cortex at two timepoints. Acute profiling was conducted at 4 min post-injection (peak head-twitch response, HTR), and chronic profiling at 12 days post-treatment. Group comparisons (DMT versus vehicle; 5-MeO-DMT versus vehicle) of metabolite peak areas were performed using two-sided Mann– Whitney U tests. Pathway enrichment analysis was conducted with MetaboAnalyst using over-representation analysis (hypergeometric test) and topology-based impact scores. Multiple-testing correction was applied using the Benjamini–Hochberg false discovery rate (FDR).

### Acute Metabolomics at Peak HTR (4 min)

At the 4 min timepoint, no individual metabolites reached statistical significance after FDR correction. Several metabolites showed nominal differences (raw P < 0.05) between psychedelic-treated groups and vehicle, including tryptophan pathway intermediates (tryptophan and serotonin). Overall, the peak HTR timepoint was not associated with robust, detectable alterations in bulk frontal cortex metabolites for either DMT or 5-MeO-DMT compared with vehicle (Supplementary Data 1).

### Long-term Metabolomics at 12 Days Post-treatment

At the 12-day timepoint, metabolites that were upregulated or downregulated within the DMT-treated group included increases in malate, adenosine, methylthioadenosine and several carnitine species (all P < 0.05) (Figure 7a). A full list of raw metabolite changes in the DMT-treated group can be found in Supplementary Data 2. DMT treatment was associated with a significant reduction in oxidized glutathione (GSSG; log₂ fold change ≈ −0.95, P = 0.003, Mann–Whitney U test; n = 8–9 per group) (Figure 7b). Pathway analysis (MetaboAnalyst) identified glutathione metabolism as the most strongly enriched pathway in the DMT-treated group (raw P = 0.002; FDR = 0.13) (Figure 7c). In the 5-MeO-DMT-treated group, enrichment of glutathione metabolism was substantially weaker (raw P = 0.070). A full list of raw metabolite changes in the 5-MeO-DMT-treated group can be found in Supplementary Data 3.

**Figure 7.**
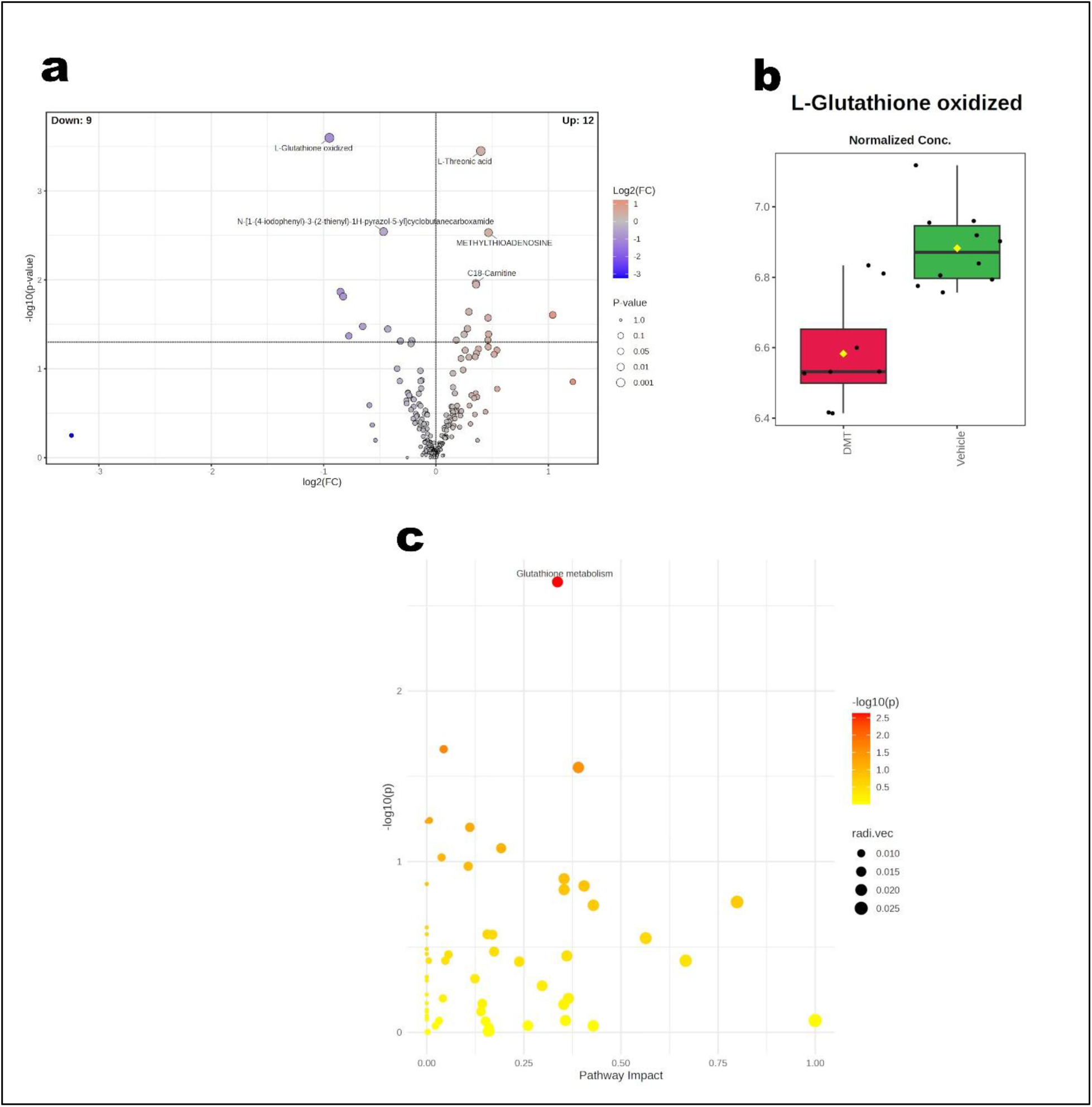
Long-term metabolomic changes 12 days after a single dose of DMT. (a) Volcano plot of metabolite changes in the DMT-treated group versus vehicle. X-axis shows log₂ fold change; y-axis shows −log₁₀(P-value). Selected significantly altered metabolites are labelled. Oxidized glutathione (GSSG) was the most statistically significant downregulated metabolite. (b) Normalized concentration of oxidized glutathione (GSSG) in the DMT and vehicle groups. DMT significantly reduced GSSG (log₂ fold change ≈ −0.95, P = 0.003, Mann–Whitney U test; n = 8–9 per group). (c) Pathway impact analysis (MetaboAnalyst) of the DMT-treated group. Glutathione metabolism was the most strongly enriched pathway (raw P = 0.002; FDR = 0.13). Group comparisons used two-sided Mann–Whitney U tests. Pathway analysis used over-representation analysis with Benjamini– Hochberg FDR correction.

Together, these data indicate that acute psychedelic effects (peak HTR) produce minimal detectable changes in bulk cortical metabolites, whereas sustained adaptations, particularly redox shifts in the DMT-treated group, are evident by day 12.

## Discussion

The present study provides a comprehensive preclinical comparison of the acute receptor pharmacology, behavioral effects, and longer-term neuroplastic consequences of the tryptamine psychedelics *N*, *N*-dimethyltryptamine (DMT) and 5-methoxy-*N*, *N*-dimethyltryptamine (5-MeO-DMT). We demonstrate that these structurally related compounds produce qualitatively distinct dose–response profiles in the head-twitch response (HTR) assay, engage both overlapping and receptor-specific mechanisms in pharmacological challenge experiments, and induce lasting increases in synaptic-plasticity-associated proteins alongside region-specific TrkB activation and metabolomic reprogramming. Critically, we show that several pharmacological strategies can attenuate HTR (a rodent proxy for hallucinogenic potential) while largely preserving the reduction in marble-burying behavior, an accepted screening assay for compounds with potential obsessive-compulsive disorder (OCD)-like therapeutic activity. These findings carry direct translational implications for the development of more tolerable and scalable psychedelic-based therapies.

Prior work has characterized the HTR-inducing effects of both DMT and 5-MeO-DMT. DMT was included in a large-scale screen of 41 hallucinogens in C57BL/6J mice and demonstrated a strong correlation between mouse HTR potency and reported human hallucinogenic potency; however, DMT effects were noted to be strain-dependent and sometimes modest ^40^. Jeferson et al.^41^, using a magnetometer-based detection method comparable to ours, recently reported that 5-MeO-DMT induces a robust but notably brief HTR (full-width at half-maximum ≈ 3–5 min versus ≈14 min for psilocybin) with high inter-animal variability in the same mouse strain. Our head-to-head dose–response data (1.25–80 mg/kg) extend these findings by revealing a clear bell-shaped profile for DMT versus a monotonic increase for 5-MeO-DMT (Figure 2).

To our knowledge, this is the first study to examine DMT and 5-MeO-DMT in the marble-burying test (MBT) and to test whether receptor modulators that blunt HTR also affect this OCD-relevant behavioral output. Previous MBT work with psychedelics has focused primarily on psilocybin ^37,42^, where 5-HT1A receptor tone was shown to be important. The dissociation we observe (attenuation of psychedelic-induced HTR by 5-HT2A antagonism, 5-HT1A agonism, or 5-HT1D antagonism without attenuation of DMT-induced marble buried reduction, and inducing marble buried reduction as a co-treatment with 5-MeO-DMT) highlights a potential pharmacological window for separating the acute, limiting subjective effects of psychedelics from some of their potential therapeutic-like behavioral actions. This is particularly relevant for OCD, where current first-line treatments are only partially effective, and psychedelic-assisted approaches are gaining clinical interest ^43,44^.

The HTR remains one of the most reliable and widely used functional readouts of 5-HT2A receptor engagement by serotonergic psychedelics ^45,46^. Our pharmacological dissection revealed both expected and novel receptor contributions to DMT and 5-MeO-DMT-induced HTR (Figure 3). As anticipated and previously demonstrated with psilocybin ^39^, the 5-HT2A antagonist M107900 (0.5 mg/kg) robustly attenuated HTR for both compounds, consistent with the established role of the 5-HT2A receptor as the primary mediator of the HTR across serotonergic psychedelics. Unexpectedly, we observed that the selective 5-HT1D antagonist LY310762 (1 mg/kg) induced near-complete suppression of HTR with DMT, 5-MeO-DMT, and psilocybin. To our knowledge, this is the first demonstration in the psychedelic literature that 5-HT1D receptor antagonism can functionally oppose the acute hallucinogenic-like effects of tryptaminergic psychedelics.

This dissociation raises the possibility that 5-HT1D receptors exert a previously under-appreciated or even independent modulatory influence on acute psychedelic-like behavioral output. While 5-HT2A receptor activation is widely viewed as the primary driver of the HTR and many subjective effects of classic psychedelics, the magnitude of HTR suppression produced by 5-HT1D antagonism was comparable to that seen with 5-HT2A blockade. This observation suggests that 5-HT1D signaling may itself function as a critical regulatory node, potentially acting as a parallel or alternative “switch” of the acute behavioral response to psychedelics, which the full neural mechanism has yet to be elucidated. Importantly, 5-HT1D receptor modulation affected HTR without consistently altering the reduction in marble-burying behavior with DMT treatment, indicating that these two effects are at least partially dissociable (Figure 4b). 5-MeO-DMT did not reduce marble burying by itself, however co-treatment of 5-MeO-DMT with 5-HT1D antagonism did produce a significant reduction in marble burying (Figure 4c). We speculate that this finding brings on a double purpose, where on one hand we reduced the HTR induced by 5-MeO-DMT with 5-HT1D antagonism, and on the other hand blocked available 5-HT1D receptors from 5-MeO-DMT binding therefore allowing more 5-MeO-DMT molecules to bind to other receptors that induce the reduction in marble burying behavior. 5-MeO-DMT exhibits substantially higher affinity for the 5-HT1D receptor (Ki ≈ 2–6 nM) relative to the 5-HT2A receptor (Ki ≈ 900–2000 nM), whereas DMT shows more balanced affinities at the two sites (Ki ≈ 39 nM at 5-HT1D and ≈ 127 nM at 5-HT2A) ^47,48^. These findings position 5-HT1D as a promising pharmacological target for attenuating acute hallucinogenic-like effects while preserving potential therapeutic-like actions, and they highlight the need for future work on 5-HT1D expression, signaling interactions with 5-HT2A, and circuit-level contributions in relevant brain regions.

A parallel and complementary finding was the significant potentiation of HTR by the 5-HT1A antagonist NAD-299 (3 mg/kg) (Figure 3). This result is the mirror image of the robust HTR attenuation we observed with the 5-HT1A agonist 8-OHDPAT (1 mg/kg) which is consistent with the well-established inhibitory role of 5-HT1A receptor tone on HTR across multiple tryptaminergic psychedelics, including psilocybin-induced HTR ^39^. The convergence with human data is particularly striking: Strassman and colleagues reported that pretreatment with the 5-HT1A antagonist/β-blocker pindolol significantly enhanced the subjective intensity and duration of intravenous DMT effects in healthy volunteers ^11^. The convergence between our rodent HTR data and Strassman’s human findings strengthens the translational relevance of the HTR assay as a proxy for certain aspects of the human psychedelic experience, at least with respect to 5-HT1A receptor tone.

The dose–response profiles of DMT and 5-MeO-DMT in the HTR assay were qualitatively distinct (Figure 2). DMT induced a clear bell-shaped curve, with significant HTR at low-to-moderate doses (1.25–5 mg/kg) that declined at higher doses (10–80 mg/kg). In contrast, 5-MeO-DMT produced no HTR at lower doses (1.25 – 2.5 mg/kg) and a robust, monotonic increase in HTR from 5 mg/kg upward. This pattern for DMT is reminiscent of the well-described transition in human phenomenology: lower doses typically elicit prominent external, open-eye visual hallucinations while the subject remains in a relatively wakeful, reality-oriented state, whereas higher “breakthrough” doses shift the experience toward intense, predominantly internal, closed-eye visionary phenomena accompanied by profound interactions with archetypical entities and loss of ordinary waking orientation ^14,49^.

Although 5-HT2A (and, as we show here, 5-HT1D) antagonists completely abolished DMT-induced HTR in mice (Figure 3a), the extent to which these receptors mediate the full subjective DMT experience remains incompletely resolved. The only published human study ^50^ examining 5-HT2A blockade of a DMT-containing preparation used ayahuasca, which produces a milder, more externally visual, and pharmacologically complex experience that rarely reaches some characteristics of the “breakthrough” state of pure inhaled or injected DMT. It is therefore plausible that ketanserin primarily attenuated the lower-dose, more serotonin-system-dominant component of the ayahuasca experience that is correlated with HTR, without testing the deeper, internal “breakthrough” phenomenology that pure DMT can rapidly induce. We speculate that HTR may preferentially capture this earlier, sensorimotor or 5-HT2A/5-HT1D-linked component of the DMT experience. At higher doses, the experience may engage additional neural substrates that are less dependent on the circuits generating head-twitch motor output, resulting in the observed decline in HTR. This interpretation is consistent with the observation that DMT produces the lowest peak HTR intensity (approximately 3–5 head twitches per peak 2 min time bin) among the tryptamines examined in similarly designed experiments, in contrast to the substantially higher peak HTR for 5-MeO-DMT (15–30), and as previously reported psilocybin (12–20), and 5-HTP (7–11) ^39^.

By comparison, the subjective effects of psilocin, 5-MeO-DMT, and LSD in humans tend to retain a more sustained open-eye visual component even at strong doses; only at very high doses do people report true ego loss or a form of disorienting, relatively weak out-of-body experience that remains substantially milder than a classic DMT “breakthrough”. While HTR is certainly an imperfect and incomplete proxy for the full human psychedelic experience, the dissociation observed here with DMT offers a useful window into which aspects of the psychedelic state the HTR may best capture. In particular, this finding suggests that HTR may primarily measure the external sensory/visual hallucination component rather than changes in cognition or consciousness. Pure DMT, even at “breakthrough” doses, is notable for preserving a high degree of cognitive clarity and self-awareness (subjects know who they are, where they are, and that they have taken a substance), in contrast to other classic psychedelics and to serotonin syndrome, both of which more readily produce profound distortions of cognition, self, identity, and orientation. If this interpretation is correct, HTR may serve as a selective preclinical screen for the visual-hallucinatory aspect of serotonin-system-dominant activation. This could prove valuable for the development of more accessible therapeutic tools that retain desired plasticity-related effects while minimizing the external visual component. It also raises the possibility that the bell-shaped HTR curve of DMT reflects a transition from a more “serotonin-like” mode of action at lower doses to a more distinctly DMT-system-dominated mode at higher doses, a distinction that may be relevant both to its endogenous role and to its unique phenomenological profile among the classic tryptamines.

Recent pathway-selective profiling at the 5-HT2A receptor has provided a useful mechanistic frame for the distinct HTR profile of DMT. In a comprehensive comparison of second-messenger and transducer pathways, Rudin et al. found that the majority of serotonergic psychedelics, as well as serotonin itself, display a significant bias toward PLC-IP1 formation (the canonical Gq–PLC readout) relative to β-arrestin2 recruitment ^51^. DMT stood out as a clear exception: it was the only compound among those tested that favored β-arrestin2 over PLC-IP1, while also showing a relative preference for the PLA2–AA pathway over PLC-IP1 ^51^. This is noteworthy because the magnitude of the head-twitch response has been shown to track 5-HT2A–Gq efficacy, with a threshold level of Gq–PLC signaling required to elicit robust HTR; β-arrestin-biased ligands generally lack this behavioral effect ^52^. The structural distinction between DMT and the other tryptamines examined is consistent with this divergence. As illustrated in Figure 1, classic tryptamine psychedelics such as psilocin, 5-MeO-DMT and bufotenine, like serotonin itself, carry an oxygen-based substituent on the indole ring, whereas DMT lacks this motif entirely. Although direct high-resolution comparisons of ligand-induced conformational ensembles are still emerging, recent cryo-EM structures of 5-HT2A bound to serotonin, psilocin and DMT show that the 5-hydroxy group of serotonin sits within hydrogen-bonding range of a key residue in the orthosteric binding pocket, an interaction that is absent in DMT (which lacks any oxygen motif) and exhibits a corresponding positional shift ^53^. One plausible interpretation is that the absence of the oxygen motif contributes to the relative under-engagement of the Gq–PLC pathway that appears to underlie the markedly lower peak HTR and inverted-U dose–response of DMT. This differential engagement may reflect a deeper physiological reality. DMT is an endogenous ligand ^3,4^, and the receptors it shares with serotonin may therefore function, under normal conditions, as a dual-input system rather than as dedicated “5-HT receptors”. A well-known parallel exists in the adrenergic system, where the two endogenous catecholamines epinephrine and norepinephrine act on the same α-and β-adrenergic receptor family yet produce distinct physiological outcomes because of differences in affinity, efficacy and tissue-specific signaling bias. If serotonin and DMT likewise engage overlapping receptors with pathway-selective consequences, then the conventional nomenclature that frames DMT simply as a “serotonergic psychedelic” may be historically skewed: the receptors in question appear to be shared resources that the organism can use according to physiological need. Viewed in this light, the compounds that possess both a dimethyltryptamine (DMT-like) core and an oxygen-containing (serotonin-like) motif can be understood as hybrid molecules (psilocin [4-OH-DMT], 5-MeO-DMT, bufotenine [5-OH-DMT], etc.) that simultaneously drive strong Gq–PLC-linked sensorimotor and visual effects that resemble those of serotonin excess (as seen in serotonin syndrome), while also engaging the plasticity-promoting and subjectively “connective” or spiritually resonant dimensions more characteristically associated with DMT. Serotonin syndrome, by contrast, is not typically reported to produce the same sense of ontological oneness or collective relatedness, and serotonin syndrome first line treatments are 5-HT2A antagonists ^54^. The low and bell-shaped HTR of pure DMT, together with its robust long-term synaptic and metabolic plasticity signature, is therefore consistent with a relative enrichment of the DMT-like contribution and a relative low activation of the pure serotonin-like sensorimotor component. As will be discussed below, this interpretation is supported by the robust long-term synaptic and metabolic plasticity signature observed after DMT. Differentiating these two contributions, both to acute subjective phenomenology and to the lasting neuroplastic changes that appear therapeutically relevant, will be essential for a more precise mechanistic understanding of tryptamine psychedelic-based treatment development that focus on the desired plasticity and spiritual effects while minimizing the accessibility-limiting acute hallucinatory experience.

In this study, our pharmacological modulations have produced several interesting results. Firstly, we observed that attenuation of HTR induced by DMT and 5-MeO-DMT treatments was confirmed for 5-HT1A agonism and 5-HT2A antagonism and extended to more novel receptors such as 5-HT1D antagonism, and in the case of DMT to 5-HT1B agonism as well (Figure 3). Secondly, we have demonstrated that co-treatment with the receptor modulation of 5-HT1A agonism, 5-HT2A antagonism and 5-HT1D antagonism that reduced the HTR did not abolish the reduction in marble burying behavior that DMT and as a 5-MeO-DMT co-treatment enhancer for reducing marble burying (Figure 4).

These pharmacological findings have direct implications for the development of more accessible psychedelic-based treatments. A major practical limitation of current psychedelic therapies is the intensity and duration of the acute subjective effects, which can limit tolerability, require extensive clinical supervision, and reduce scalability. Our data indicates that it is possible, at least in preclinical screening assays, to attenuate HTR (a proxy for hallucinogenic potential) through targeted receptor modulation (notably at 5-HT1D, 5-HT2A, and 5-HT1A) while largely preserving/enabling the reduction in marble-burying behavior produced by the psychedelic itself. The marble-burying test is an accepted screening assay for compounds that alleviate OCD-like symptoms; several clinically used agents, including 5-HT1A agonists such as buspirone, reduce burying in this paradigm ^37^. The observation that certain co-treatments can blunt HTR without eliminating the MBT effect supports the feasibility of combination strategies aimed at reducing the acute “trip” while retaining therapeutic-like behavioral activity. Whether such pharmacological tuning can be translated to humans, and whether it preserves other clinically relevant outcomes, remains an important question for future investigation.

Additionally, our data revealed a differential involvement of the 5-HT1B receptor. The selective 5-HT1B agonist CP 94253 (5 mg/kg) significantly attenuated DMT-induced HTR and completely prevented the reduction in marble burying produced by DMT (Figures 3, 4), demonstrating the involvement of the 5-HT1B receptor in OCD-like symptom-alleviating behavioral effects of DMT in this assay. In striking contrast, while 5-HT1B agonism produced only a non-significant trend toward HTR reduction with 5-MeO-DMT, the combination proved lethal in MBT, resulting in mortality that precluded behavioral testing. It should be noted that the HTR experiment used younger (11-13 weeks old, ∼25-30g) C57BL mice, and the mortality was noted in the MBT that used older (12-26 weeks old∼30-50g) ICR mice. This mortality likely stems from a pharmacodynamic interaction: 5-HT1B agonists induce cranial and systemic vasoconstriction and are clinically utilized for acute migraine relief precisely through this mechanism of blood vessel contraction ^55^. 5-MeO-DMT, in turn, is well-documented to elicit robust increases in heart rate ^56^. We propose that the conjunction of pronounced tachycardia with intense vasoconstriction may precipitate acute circulatory failure. This observation raises a critical safety concern for potential human applications involving the co-administration of 5-MeO-DMT with 5-HT1B agonists or other vasoconstrictive agents.

On the molecular level, acute administration of both DMT and 5-MeO-DMT increased TrkB phosphorylation (p-TrkB/TrkB ratio) in a region-and compound-specific manner 1h after injection, as measured by immunofluorescence (Figure 5a). DMT induced more widespread and robust elevations across the default-mode network and hippocampal regions, whereas 5-MeO-DMT effects were more restricted (primarily the frontal association cortex, and prelimbic cortex). At the same 1h post DMT-injected time point, automated capillary western blotting (ABBY) revealed no significant increase in total BDNF or TrkB protein levels in the hippocampus (Figure 5b-e). This dissociation is noteworthy when viewed against the seminal work of Moliner et al. ^35^, who showed that LSD and psilocin bind directly to the transmembrane domain of TrkB dimers with nanomolar affinity and function as positive allosteric modulators that potentiate and prolong BDNF-induced TrkB dimerization and downstream signaling in a manner independent of 5-HT2A receptor activation. In contrast, other proposals in the field have emphasized 5-HT2A-dependent mechanisms for psychedelic-induced plasticity. Vargas et al. ^57^, for example, highlighted the contribution of intracellular 5-HT2A receptor populations to neuroplasticity and enhanced BDNF signaling without reference to direct psychedelic TrkB allosteric binding, while models such as those presented by Werle et al. ^24^ propose that 5-HT2A activation is the main driver that promotes BDNF release or expression, which in turn drives TrkB signaling. Our finding of clear p-TrkB elevation without a detectable increase in BDNF protein 1 hour post psychedelic injection aligns more closely with the direct allosteric facilitation model of Moliner et al. than with mechanisms that require acute upregulation or release of BDNF secondary to 5-HT2A stimulation. Our data are consistent with a substantial 5-HT2A-independent component, although definitive separation of the pathways will require targeted follow-up studies (e.g., 5-HT2A/TrkB antagonism). Whether the observed increase in p-TrkB leads to sustained engagement of canonical downstream effectors such as AKT and mTOR also remains to be determined.

A particularly intriguing and translationally relevant finding was the significant reduction in hippocampal BDNF protein levels observed 12 days after a single dose of DMT (5 mg/kg), with no comparable change after 5-MeO-DMT (Figure 5d). This long-term downregulation occurred in the context of robust acute TrkB phosphorylation and subsequent increases in multiple synaptic-plasticity-related proteins. One plausible interpretation is homeostatic regulation: strong, widespread activation of the TrkB system by exogenous DMT may trigger compensatory mechanisms that reduce BDNF production in the ensuing days to weeks, thereby preventing sustained overstimulation. Two non-mutually exclusive scenarios can be envisioned. In one, the compensatory reduction overshoots and temporarily deprives the system of an optimal BDNF tone (a potentially maladaptive outcome). In the other, the brain, having experienced a period of heightened TrkB signaling, downregulates BDNF production because less ligand is required to maintain the desired level of pathway activity, an energy-efficient optimization. Distinguishing these possibilities, determining the time course of BDNF recovery, and testing whether lower, but still-plasticity-effective doses of DMT can achieve strong acute p-TrkB activation without the later BDNF reduction are important directions for future work. Because the DMT 5 mg/kg dose used here corresponds to a translationally relevant dosing in ongoing clinical investigations of DMT, any therapeutic development targeting the TrkB pathway with DMT or related compounds will need to carefully map these longer-term dynamics to ensure net benefit rather than unintended negative homeostatic disruption.

Despite the nuanced BDNF response, both DMT and 5-MeO-DMT produced significant long-term (12-day) increases in several synaptic-plasticity-associated proteins (PSD95, synaptophysin, and for DMT, SV2A as well) across frontal cortex, amygdala, and hippocampus (Figure 6), with effects comparable in direction and magnitude to those previously reported with psilocybin and psychedelic mushroom extract ^36^. These proteins are established markers of structural and functional synaptic remodeling ^58,59^. Several of these proteins have been linked in the literature to TrkB signaling and BDNF-mediated plasticity, providing a coherent molecular bridge between the acute p-TrkB activation we observed, and the enduring protein changes measured at 12 days. The three major cellular pathways initiated by TrkB signaling upon BDNF binding are the MAPK/ERK (Ras/Raf/MEK/ERK), PI3K/Akt, and PLC-γ cascades, which collectively drive neuronal survival, differentiation, gene expression changes, and synaptic plasticity through both transcriptional and local signaling mechanisms ^60^. PSD-95 expression and postsynaptic localization are increased by all three TrkB signaling cascades ^61^. Synaptophysin, a core presynaptic vesicle marker, is elevated by BDNF/TrkB signaling with clear contributions from the MAPK/ERK and PI3K/Akt pathways ^62^. The pattern is consistent with the broader hypothesis that tryptamine psychedelics can initiate a cascade of neuroplastic adaptations that persist well beyond the acute pharmacological effects.

The untargeted metabolomic analysis complements these protein findings. At 4 min post-injection (near the peak of the head-twitch response), no individual metabolites reached statistical significance after rigorous filtering, indicating that the acute behavioral phenotype is not accompanied by rapid, large-scale shifts in the frontal cortical metabolome detectable by this platform. In contrast, at the 12-day time point we observed significant alterations, most prominently after DMT treatment. These included a significant reduction in oxidized glutathione (GSSG) and enrichment of the glutathione metabolism pathway, together with increases in malate, adenosine, adenine, methylthioadenosine, and several acylcarnitines (Figure 7). A more reduced glutathione redox state has been shown to support the maintenance of synaptic plasticity mechanisms ^63^. Concurrently, elevated TCA-cycle intermediates (malate), purine metabolites (adenosine, adenine, MTA), and acylcarnitines are consistent with enhanced mitochondrial energy metabolism and processes that supply the high energetic and biosynthetic demands required to sustain elevated levels of synaptic proteins ^64,65^. In contrast, 5-MeO-DMT produced a substantially weaker enrichment of glutathione metabolism (raw P = 0.070) or elevated energy-and purine-related metabolites at the 12-day time point. This suggests either that any plasticity-supporting metabolic adaptations induced by 5-MeO-DMT had already subsided by day 12, or that the long-term metabolic programs associated with sustained synaptic remodeling are not fully shared between the two compounds.

Taken together, the glutathione and energy-related metabolic signatures at day 12 are most consistent with a plasticity-maintenance phase, in which the tissue has adapted to support the structural changes already in place (increased synaptophysin, PSD-95, and SV2A). The absence of stronger inductive metabolic signatures at this late time point raises the possibility that earlier sampling windows (e.g., 1–6 days post-administration) would be better suited to capture the peak metabolic programs that actively drive the observed long-term increases in synaptic proteins.

Overall, the present work provides a broad preclinical characterization of DMT and 5-MeO-DMT that spans acute receptor pharmacology, a clinically relevant behavioral screening assay (marble burying as a test for potential alleviation of OCD-like symptoms), acute TrkB activation, and long-term molecular and metabolic plasticity markers. The identification of 5-HT1D receptor antagonism as a potent suppressor of HTR, without apparent loss of the MBT effect for DMT and as a reducer of marble burying behavior as a co-treatment with 5-MeO-DMT, represents a novel pharmacological lead that could inform strategies to improve the tolerability and accessibility of psychedelic-based therapies. At the same time, the TrkB and downstream protein data reinforce the emerging view that direct allosteric modulation of TrkB, in addition to or instead of classical 5-HT2A signaling, contributes to psychedelic-induced neuroplasticity, while also revealing previously unappreciated longer-term homeostatic regulation of BDNF itself after exogenous DMT treatment. These findings underscore both the promise and the complexity of harnessing DMT and 5-MeO-DMT for neuropsychiatric applications and highlight specific receptor targets and dose considerations that warrant further translational attention. More broadly, the distinct behavioral and molecular signatures of DMT observed here are consistent with the possibility that receptors traditionally viewed as serotonergic may innately function, at least in part, as shared resources for more than one endogenous ligand, and that differentiating the respective contributions of serotonin-like and DMT-like signaling will be important for future mechanism-guided development of tryptamine psychedelic-based therapeutics and understanding.

## Methods

### Animals

All experiments were performed on adult (11-13 weeks old, ∼25-30g) C57BL/6J male mice, apart from the marble burying test (MBT) which was conducted on male 12-26 weeks old ICR mice that weighed ∼30-50g. Sample size was based on prior data. Animals were housed under standardized conditions with a 12-h light/dark cycle, stable temperature (22 ± 1 °C), controlled humidity (55 ± 10%) and free access to food and water. Mice were assigned to experimental groups by randomly extracting them from the holding cages. All experiments were conducted with the investigator blind to treatment assignment. Experiments were conducted in accordance with AAALAC guidelines and were approved by the Authority for Biological and Biomedical Models Hebrew University of Jerusalem, Israel, Animal Care and Use Committee # MD-21-16563-4. All efforts were made to minimize animal suffering and the number of animals used.

### Drugs

N,N-DMT (purity=99%) was supplied by Lucy Scientific Design, Canada. 5-MeO-DMT (purity =99%) was supplied by Nextar, Israel. Psilocybin (purity=99%) was supplied by Usona Institute, Madison, WI, USA. N,N-DMT, and 5-MeO-DMT were administered by i.p. injection immediately before the assessment of HTR at the dose of 1.25-80 mg/kg, or at 5 mg/kg (N,N-DMT) and 10 mg/kg i.p (5-MeO-DMT) preceded by the 5-HT2A antagonist M107900 (0.5 mg/kg), the 5-HT1A agonist 8-OHDPAT (1 mg/kg), 5-HT1A antagonist NAD-299 (3 mg/kg), 5-HT1B agonist CP 94253 (5 mg/kg), 5-HT1B antagonist SB 224289 (2.5 mg/kg), 5-HT2C agonist WAY-161503 (3 mg/kg), 5-HT2C antagonist RS-102221 (4 mg/kg), 5-HT1D agonist PNU 142633 (0.3 mg/kg), 5-HT1D antagonist LY310762 (1 mg/kg), 5-HT7 agonist AS-19 (20 mg/kg), 5-HT7 antagonist SB-269970 (60mg/kg), TAAR1 antagonist EPPTB (10mg/kg), and σ receptor antagonist BD-1047 (10 mg/kg). Psilocybin was injected at a dose of 4.4mg/kg preceded by the 5-HT1D agonist PNU 142633 (0.3 mg/kg), and 5-HT1D antagonist LY310762 (1 mg/kg). In all cases, injections were administered in a standard injection volume of 10 µl/g per mouse. N,N-DMT, 5-MeO-DMT, and psilocybin were rapidly dissolved completely in saline (0.9% NaCl). Control mice received saline injections (Vehicle – VEH).

### Head Twitch Response

Head twitch response (HTR) was measured over 20 minutes by means of a magnetometer apparatus as described by Shahar et al. ^39^. Briefly, small neodymium magnets (N50, 3mm diameter × 1mm height, 50 mg), were attached to the outer ears of mice. After a 5–7-day recovery period, the ear-tagged animals were placed inside a magnetometer apparatus (supplied by Mario de la Fuente Revenga PhD. of Virginia Commonwealth University) immediately after injection of the test compounds for 20 min of measurement (Supplementary Video 1). The output was amplified (Pyle PP444 phono amplifier) and recorded at 1000 Hz using a NI USB-6001 (National Instruments, US) data acquisition system. Recordings were performed using a MATLAB driver (MathWorks, US, R2021a version, along with the NI myDAQ support package) with the corresponding National Instruments support package for further processing. A custom MATLAB script was used to record the processed signal, which was presented as graphs showing the change in current as peaks (mAh). A custom graphical user interface created in our laboratory was used to further analyze the recording into an Excel spreadsheet.

## Marble-burying test (MBT)

MBT (Supplementary Video 2) was performed in transparent cages containing ∼4.5 cm fine sawdust, as previously described ^37,66^. Twenty glass marbles were placed equidistant from each other in a 5 × 4 pattern. The experiment was done under dim light in a quiet room to reduce the influence of anxiety on behavior. The mice were left in the cage with the marbles for a 30-min period after which the test was terminated by removing the mice. A marble was considered buried when two thirds or more of its size was covered with burying substrate and number of buried marbles was counted after 30 minutes (Supplementary Video 2). All mice underwent a pretest without any injection and the number of marbles buried was counted. Only mice that buried at least 15 marbles were selected to perform the test after drug administration. 80% of pretested mice fulfilled this criterion and were used in the definitive experiment, which took place at least a week following the pretest.

### Euthanasia and tissue collection

For Western blot and metabolomics analyses, mice were placed in a sealed 2 L induction chamber equipped with a perforated false floor that occupied the lower quarter of the chamber volume. Cotton balls were placed beneath the perforated floor, and approximately 1 mL of liquid isoflurane was applied to the cotton balls every 7 minutes. This setup allowed controlled vaporization of isoflurane while preventing direct contact between the mice and the liquid anesthetic, thereby avoiding skin irritation. Anesthesia depth was confirmed by loss of the righting reflex and absence of a response to paw pinch. Once surgical anesthesia was achieved, mice were euthanized by cervical dislocation, and tissues were rapidly collected for protein extraction and metabolomic analysis.

For immunofluorescence studies, 1h post i.p. injection of either vehicle, DMT (5 mg/kg), or 5-MeO-DMT (5 mg/kg) mice were deeply anesthetized with isoflurane (delivered via a precision vaporizer and nose-cone mask system at an induction concentration of 3–4% in oxygen or medical air/oxygen mixture, maintained at 1–2% as needed to sustain a surgical plane of anesthesia, confirmed by loss of righting reflex and absence of response to toe pinch). Mice were then transcardially perfused with 5 ml of phosphate-buffered saline (PBS) containing heparin (10 U/ml) to clear blood from the vasculature, followed by 10 ml of 4% paraformaldehyde (PFA) for tissue fixation. Brains were rapidly dissected, post-fixed overnight in 4% PFA at 4°C, incubated in 30% sucrose in PBS for 48 h at 4°C, then frozen in OCT compound and processed for immunofluorescence staining.

### Immunofluorescence p-TrkB/TrkB

Sagittal brain sections 35 μm in thickness were prepared using a cryostat (Leica CM 1950 Cryostat). Sections were organized and Bregma 0.24mm were selected based on brain atlas ^67^. Sections were washed with PBS, incubated with blocking solution (PBS containing 5% donkey serum and 0.3% Triton X-100) for 1h at RT, then incubated with primary antibodies in blocking solution (goat anti-TrkB 1:100, R&D Systems, AF1494 and rabbit anti-pTrkB 1:200, Novus, NBP1-03499) at 4°C overnight, washed 3 times with PBST (PBS with 0.05% Triton X-100) and then the secondary antibodies (1:250, Alexa Fluor 488 Donkey Anti-Rabbit IgG, Jackson, 711-546-152 and 1:500, Alexa Fluor 647 Donkey Anti-Goat IgG, Jackson, 705-605-003) diluted in blocking solution were added at RT for 1h. Sections were washed once with PBST, stained with Hoechst (2μg/mL), and washed with PBST two more times. Sections were transferred onto Super Frost slides and mounted under glass coverslips with Eukitt mounting media.

Fluorescence images were acquired using a Nikon spinning disk confocal microscope (Yokogawa W1 spinning disk mounted on a Ti2E motorized inverted microscope body) equipped with 405 nm, 488 nm, and 638 nm lasers, two sCMOS ZYLA cameras, a Piezo Z stage, and NIS-Elements software for multidimensional acquisition and analysis. Large tiled (stitched) z-stack images encompassing the entire sagittal plane of each section at Bregma +0.24 mm (including the DMN and hippocampus) were captured at 20× magnification using a CFI Plan Apochromat VC 20×/0.75 NA dry objective. Z-stacks were acquired with a 1 μm step size; the number of z-planes per section ranged from 15 to 40 depending on mounting variations, with the upper and lower z-limits for each section determined by visual inspection of signal focus across all regions of interest (ROIs). The 488 nm laser was used to excite Alexa Fluor 488 (labeling pTrkB), the 638 nm laser to excite Alexa Fluor 647 (labeling TrkB), and the 405 nm laser for Hoechst.

Following acquisition, images were processed in Fiji (ImageJ) using the BigWarp plugin. A digital reference image of the Bregma +0.24 mm atlas plate was imported and non-rigidly warped to align with the morphology of each individual experimental section. This alignment enabled precise localization of brain areas on the corresponding high-resolution microscope images. ROIs for each brain region were then defined based on the warped atlas overlays. Quantitative analysis of fluorescence signal intensity was performed on the original captured z-stack images in NIS-Elements. For each section and each channel independently, signal-to-noise thresholds for pTrkB and TrkB were determined by two blinded scorers, with each scorer performing the threshold determination twice per section. The signal intensity for each signal on each section was averaged together for statistical analysis. Representative immunofluorescent image, and representative wrapped atlas image can be seen in Supplementary Figure 2 and Supplementary Figure 3 respectively.

### Western blotting (SDS-PAGE)

The frontal cortex, amygdala, and hippocampus were dissected and stored at-80°C, lysed in Pierce RIPA sample buffer (Thermo Scientific, USA), supplemented with protease inhibitor cocktail (Roche Diagnostics, Germany) and boiled for 10 min. Equivalent amounts of protein extracts (20 mg) were analyzed by SDS– 12% PAGE, followed by transfer of the proteins to polyvinylidene fluoride membrane. Blots were blocked in 5% fat free milk in TBST buffer (Tris-Tween-buffered saline) and incubated in primary antibodies, one hour at room temperature. Primary antibodies included rabbit anti-GAP43 (ab75810, 1:2000; Abcam, UK), rabbit anti-PSD95 (ab238135, 1:2000; Abcam, UK), rabbit anti-Synaptophysin (ab32127, 1:2000; Abcam, UK), rabbit anti-SV2A (ab254351, 1:1000; Abcam, UK) and mouse anti-β-Actin (8H10D10, 1:5000, Cell Signaling Technology). Blots were washed 3 times and incubated with a horseradish peroxidase-conjugated secondary antibodies (1:5000, ABclonal, China) for 1 h, followed by repeated washing with TBST buffer. Proteins were visualized by using enhanced chemiluminescence (ChemiDoc Reader MP, Bio-Rad, USA). Band intensities for each target protein were normalized to β-actin, which was used as the loading control. β-actin was selected because recent studies in the field of psychedelics and the utilization of psilocybin have employed Western blot analysis, using beta-actin exclusively as a housekeeping gene ^68–71^. Representative SDS-PAGE immunoblots can be seen in Supplementary Figure 4.

### Western Blot – ABBY

Protein expression levels of BDNF and TRKB were quantified by automated capillary western blotting using the ABBY instrument (Bio-Techne®). The antibodies utilized were anti-BDNF (ab108319, Abcam; 1:25 dilution) and anti-TRKB (TrkB Rabbit mAb #4603, Cell Signaling Technology; 1:50 dilution). Samples were prepared according to the manufacturer’s protocol. Briefly, a 5X fluorescent master mix containing 400 mM dithiothreitol and 10X sample buffer was made. The biotinylated ladder was prepared by mixing 10X sample buffer, 400 mM dithiothreitol, and deionized water. Lysates were combined with the 5X master mix to a final protein concentration of 0.3 mg/ml. Samples and ladder were heat-denatured at 95°C for 5 minutes before loading onto 12–230 kDa Separation Modules (SM-W004; Bio-Techne). Primary antibodies and luminol-S/peroxide substrate were loaded per the assay plate layout. Protein separation and immunodetection were carried out using the Anti-Rabbit Detection Module (DM-001; Bio-Techne) and Total Protein Detection Module (DM-TP01; Bio-Techne). Data were acquired using ABBY software, with total protein concentration ratios calculated in Microsoft Excel. Representative ABBY immunoblots can be seen in Supplementary Figure 5.

### Metabolomics

Sample Preparation: Frozen cortex tissues were dissected, weighed and placed in Precellys homogenization tube with beads. A ratio of 1 ml of extraction solvent per 30 mg of tissue was added. The samples were homogenized in a Precellys 24 with a Cryolys temperature controller, at approximately 0°C (3*30 sec with a 30 sec gap between cycles). The samples were then centrifuged in the Precellys tubes at 13,000 RPM for 20 min at 4°C. The supernatant was then collected into HPLC vials. All of the samples were pooled into one QC (Quality Control) sample. The samples in each treatment group were pooled into one ID (Identification) sample for MS2 (tandem mass spectrometry) purposes

LCMS Acquisition: LCMS metabolomics analysis was performed as described previously ^72^. Briefly, a Dionex Ultimate 300 high-performance liquid chromatography (UPLC) system, coupled to an Orbitrap Q-Exactive Plus Mass Spectrometer (Thermo Fisher Scientific), with a resolution of 70,000 at a 200 mass/charge ratio (m/z), electrospray ionization in the HESI source, and polarity switching mode to enable both positive and negative ions across a mass range of 70 to 1000 m/z, was used. The UPLC setup included a ZIC-pHILIC column (SeQuant; 150 mm × 2.1 mm, 5 μm; Merck) with a Sure-Guard filter (SS frit 0.5 μm). Five µL of the tissue extracts were injected, and the compounds were separated with a mobile phase gradient of 15 min, starting at 20% aqueous (20 mM ammonium carbonate adjusted to pH 9.2 with 0.1% of 25% ammonium hydroxide) and 80% organic (acetonitrile), and ending with 20% acetonitrile. The flow rate and column temperature were maintained at 0.2 mL/min and 45°C, respectively, for a total run time of 26 min. QC pool samples were run at regular intervals to monitor system stability and performance throughout the analytical batch. All metabolites were detected with mass accuracy below 5 ppm. Thermo Xcalibur software was used for data acquisition.

Metabolite Identification and Validation: Data analysis was performed using Compound Discoverer 3.3 SP2. Identification and confirmation of the exact mass of singly charged ions were performed based on their known retention times, utilizing an in-house MS library. MS2 (tandem mass spectrometry) analysis for further validation involved the fragmentation of selected ions and the analysis of the resulting fragments to gain additional structural information.

## Statistical analysis

Data are presented as mean ± SEM. Normality was assessed using the Shapiro–Wilk test. When the raw data met the assumption of normality, comparisons between two groups were performed using Welch’s unpaired t-test, and comparisons among three or more groups were performed using one-way ANOVA followed by Dunnett’s post-hoc test. For the TrkB phosphorylation experiment across multiple brain regions, data were log-transformed and analyzed by two-way ANOVA followed by Tukey’s multiple comparisons test. For synaptic protein levels measured across multiple brain regions, nested one-way ANOVA was performed followed by Dunnett’s post-hoc test with brain region nested within treatment, followed by Dunnett’s post-hoc test versus vehicle. When normality was violated on the raw data, a log transformation was applied using the equation Y = log(Y + 1). If the log-transformed data met normality assumptions, parametric tests were performed on the transformed values and results are reported on the original scale. When log transformation did not restore normality, the non-parametric Mann–Whitney U test (two groups) or Kruskal–Wallis test followed by Dunn’s post-hoc test (three or more groups) was used. All statistical analyses were conducted using GraphPad Prism version 10.2.1.

## Data Availability

The data that support the findings of this study are available from the corresponding author upon reasonable request.

## Supporting information

Shahar-Supplementary Information

## Acknowledgements

The research was supported in part by Negeb Labs. O. Shahar is supported by the President of the State of Israel Scholarship for Excellence and Scientific Innovation awarded to O. Shahar, and the Hadassah BrainLabs Center for Psychedelic Research. We thank the staff of the Hebrew University Animal Facility for excellent animal care.

## Author contributions

O. Shahar designed and performed all experiments, analyzed data, prepared figures and wrote the manuscript. A.B. contributed in co-performing and analyzing the HTR, MBT, and 12 days synaptic protein experiments as well as manuscript editing. A.S. and E.L. contributed by performing and analyzing the BDNF/TrkB ABBY Western blot experiment. P.G. and M.B.A. contributed by co-performing the MBT experiments. O. Shalev contributed by performing and analyzing the metabolomics experiment. T.L. and B.L. conceived and supervised the project, secured funding, and edited the manuscript. All authors reviewed and approved the final manuscript.

