## Supplementary material for "Preclinical Comparison of DMT and 5-MeO-DMT Reveals Behavioral Dissociation, Distinct TrkB Activation and Differential Plasticity Profiles": Shahar-Supplementary Information

Supplementary Information for:
 ***Preclinical Comparison of DMT and 5-MeO-DMT Reveals Behavioural Dissociation, Distinct TrkB Activation and Differential Plasticity Profiles*.**

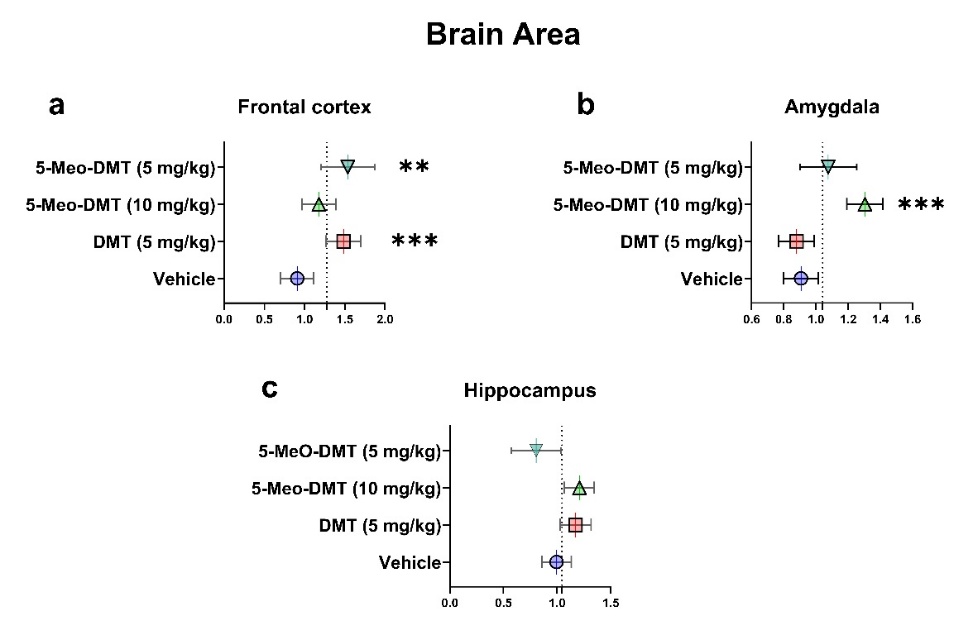

Supplementary Figure 1 - Long-term effects of a single dose of DMT or 5-MeO-DMT on synaptic proteins by brain region. Levels of GAP43, PSD95, synaptophysin, and SV2A were quantified by Western blot 12 days after treatment and analyzed by nested one-way ANOVA within each brain region. **a** Frontal cortex: significant increases following DMT (5 mg/kg) and 5-MeO-DMT (5 mg/kg) compared to vehicle. **b** Amygdala: significant increase following 5-MeO-DMT (10 mg/kg) compared to vehicle. **c** Hippocampus: no significant differences compared to vehicle for any treatment. Data are mean ± SEM normalized to β-actin. Nested one-way ANOVA followed by Dunnett’s post-hoc test. Significance vs. Vehicle; ** = p < 0.01, *** = p < 0.001. n = 5–16 per group.

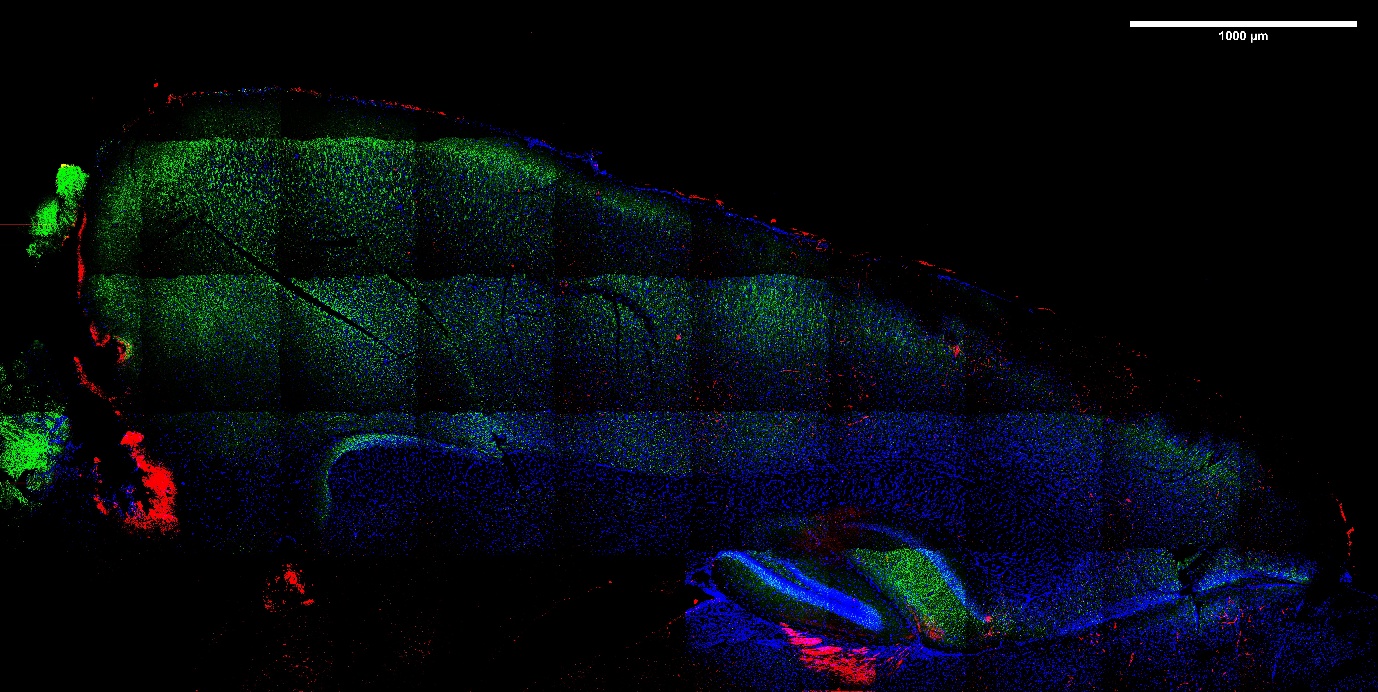

Supplementary Figure 2 – Representative sagittal immunofluorescence image of TrkB and phosphorylated TrkB (p-TrkB) following DMT administration. Sagittal brain section from a DMT-treated mouse (5 mg/kg, 1 h post-injection) showing immunofluorescence staining for p-TrkB (green), unphosphorylated TrkB (red), and nuclei (DAPI, blue). Scale bar = 1000 µm.

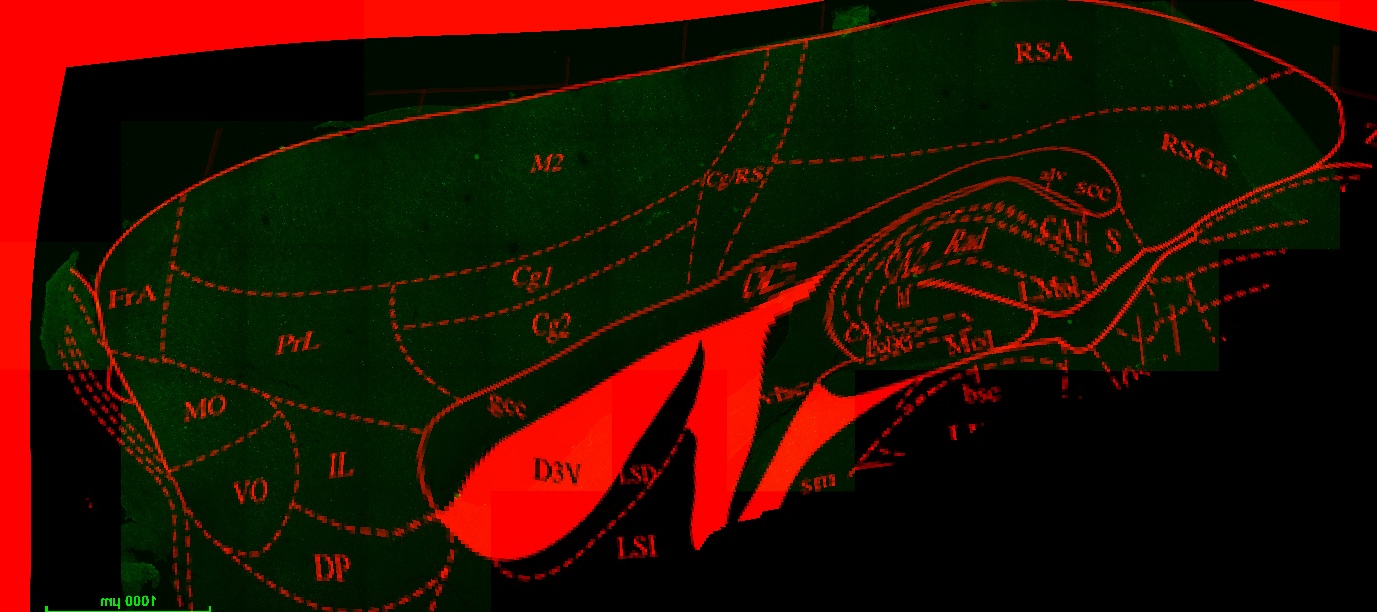

Supplementary Figure 3 – Representative image showcasing the merged wrapped atlas image overlayed on top of the specific microscope image it was wrapped to. This warpped image was used as a visual marker to draw the ROI’s of the desired brain area in NIS elements for intensity quantification.

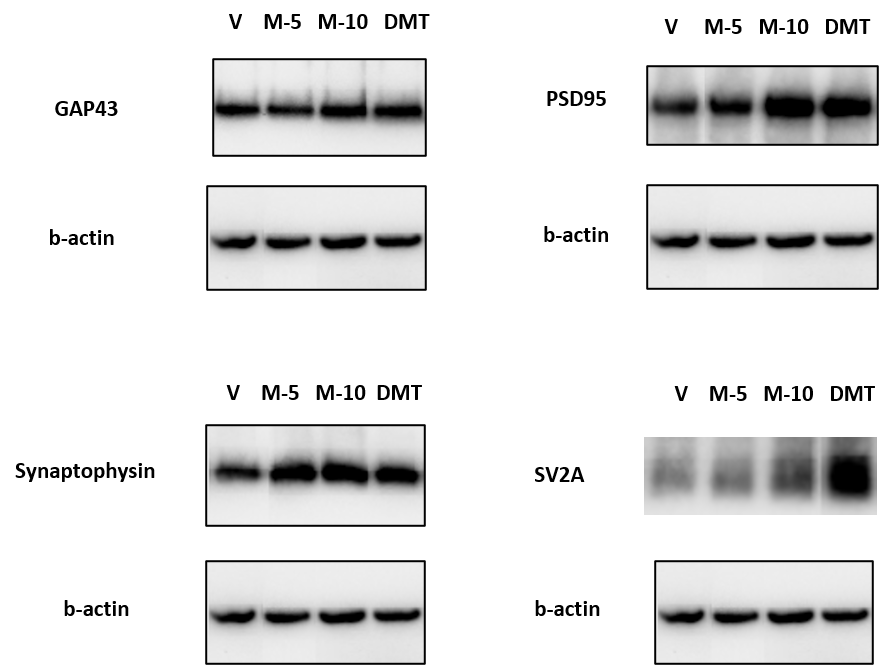

Supplementary Figure 4 - Representative immunoblots of GAP43, Synaptophysin, SV2A, PSD95 and b-actin protein expression across all brain regions after Vehicle (V), 5-MeO-DMT (5 mg/kg) (M-5), 5-MeO-DMT (10 mg/kg) (M-10), and DMT (5 mg/kg) treatments.

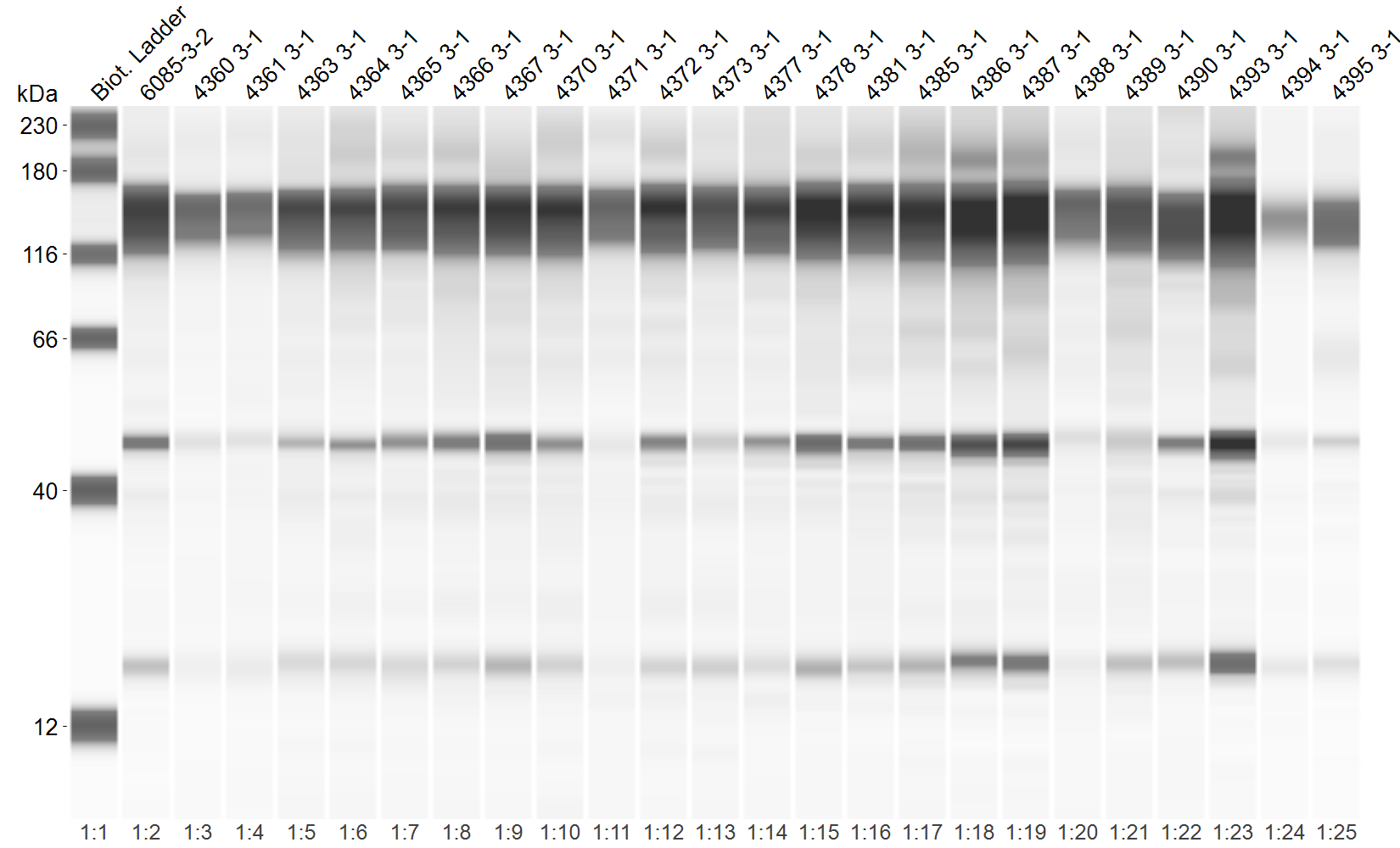

Supplementary Figure 5 - Representative automated capillary Western blot for the DMT experimental runs. The blot demonstrates the expression of TrkB (~140 kDa), mature BDNF (~20 kDa) and a putative protein aggregate (~48 kDa) across the analyzed hippocampal samples, exhibiting varying signal intensities between individual lanes. Target expression levels were quantified by integrating the area under the respective electropherogram peaks, with final expression values normalized to the total protein content measured for each individual sample.
